# HuR enhances SARS-CoV-2 non-structural protein translation through the genomic 5’-UTR, by promoting polypyrimidine tract-binding protein binding

**DOI:** 10.1101/2023.03.15.532790

**Authors:** Harsha Raheja, Risabh Sahu, Trinath Ghosh, Priya Rani, Biju George, Oyahida Khatun, Shashank Tripathi, Saumitra Das

## Abstract

Severe acute respiratory syndrome-coronavirus-2 (SARS-CoV-2) viral RNA associates with different RNA-binding host proteins at each stage of its life cycle. We found sequence dependent binding of one such important protein, human antigen R (HuR) to SARS-CoV-2 5′UTR and studied its potential role in virus life cycle. The knockdown and knockout studies revealed importance of such binding in viral translation. We identified 5′-UTR mutations in SARS-CoV-2 variants of concern that altered the HuR-binding affinity. Interestingly, HuR enhanced non-structural protein translation through the genomic 5′-UTR, by promoting polypyrimidine tract-binding protein binding to the 5′-UTR. However, HuR suppressed the structural protein translation from sub genomic 5’UTR. HuR knockout increased the sensitivity to remdesivir treatment by decreasing its half-maximal inhibitory concentration. An antisense oligonucleotide (whose binding site overlapped the HuR-binding site) reduced viral RNA levels in wild-type cells but not HuR-knockout cells. Our results indicate that HuR regulates the balance between SARS-CoV-2 structural and non-structural proteins and guides the infection of viral variants, implying that HuR can potentially be targeted for therapeutic interventions.

**Author Summary:** Viruses interacts with various host proteins throughout their life cycle. One significant protein is HuR, an RNA-binding protein that regulates RNA stability and translation. HuR binds to viral RNAs at the 5’UTR or 3’UTR, impacting their translation and replication. We identified conserved HuR binding sites in the SARS-CoV-2 5’UTR across different beta coronaviruses. This binding enhanced the initiation of translation from the genomic 5’ UTR, increasing the production of non-structural proteins essential for viral replication. Additionally, we discovered that another host protein, PTB, promotes HuR binding to the viral 5’ UTR, facilitating its loading onto ribosomes. Conversely, HuR plays an antagonistic role concerning subgenomic RNAs (sgRNAs), which code for structural proteins, by regulating and limiting their levels. This dual regulation indicates that the virus exploits HuR for its benefit while the host employs it to control viral spread. Targeting HuR may help manipulate the SARS-CoV-2 life cycle. We found that HuR knockout increased sensitivity to the antiviral drug Remdesivir. Using an antisense oligonucleotide to block HuR binding effectively reduced viral RNA levels. Our findings highlight the critical role of HuR in regulating viral protein production and its potential as a therapeutic target.

## Introduction

Severe acute respiratory syndrome-coronavirus-2 (SARS-CoV-2) is a positive-strand RNA virus with a 30-kilobase genome. The viral life cycle begins with attachment of the virus to the angiotensin-converting enzyme-2 (ACE-2) receptor. That interaction is followed by receptor-mediated endocytosis, where the virus particles are internalized into endosomes, which release viral RNA into the host cell cytoplasm. Viral RNA undergoes translation to generate a polyprotein that codes for nonstructural proteins. These non-structural proteins initiate replication on the viral 3′-untranslated region (3′-UTR), generating negative-strand genomic RNA. The positive sense genomic RNA is also used as a template to synthesize negative sense sub-genomic RNAs, which are transcribed to positive sense sub-genomic RNAs coding for viral structural and accessory proteins. The structural proteins thus formed coat the viral positive-strand RNA to form infectious virions, which are released from cells [1].

SARS-CoV-2 RNA and proteins interact with several host proteins to modulate viral pathogenesis. The viral RNA possesses binding sites for cellular RNA-binding proteins (RBPs), which assist the above-mentioned processes at different levels [2, 3]. The viral coding region is flanked by UTRs at both the 5′ and 3′ ends, which provide scaffolds for viral and cellular RBPs to bind and regulate translation, replication, or packaging. A dynamic association exists between RBPs and viral RNA during different stages of viral infection. High-throughput screens have identified many RBPs, such as IMP-1, DDX-1, and hnRNPs, that bind to SARS-CoV-2 genomic RNA [4, 5]. Here, we studied RBPs predicted to bind to the viral 5′-UTR. Of these, we selected human antigen R (HuR; also known as ELAVL1) for further study.

HuR is an RBP that binds to AU-rich elements in the 3′-UTRs of target mRNAs and stabilizes mRNAs [6, 7]. HuR shuttles between the nucleus and cytoplasm and participates in the cellular transport of various RNAs. Different post-translation modifications of HuR have been linked to different functional roles and subcellular-localization sites in cells [8]. HuR can regulate the replication of hepatitis C virus [9, 10], coxsackievirus B3 [11], and adenovirus [12]; stabilize Sindbis virus RNA [13]; and modulate human immunodeficiency virus reverse transcription [14]. It was also shown to bind the SARS-CoV-2 genome in genome-wide screens [4, 15]. We confirmed that HuR can bind the viral 5′-UTR through multiple independent assays and studied its importance in the SARS-CoV-2 life cycle. We examined the potential roles of HuR in regulating the translation of both structural and non-structural proteins, which are translated from separate sub-genomic and genomic RNAs, respectively. Further, we blocked HuR–SARS-CoV-2 genome interactions with specific antisense oligonucleotides (ASOs), suggesting that they can be explored as therapeutics for treating COVID19 disease.

## Results

### HuR regulated the SARS-CoV-2 life cycle

The 5′-UTR of SARS-CoV-2 is 265 nucleotides (nt) long. Enzymatic probing showed that the initial 300 nt of the SARS-CoV-2 genome at the 5′ end forms five stem-loop secondary structures that are conserved across all emerging variants[16]. The RNA-binding protein database[17] (RBPDB) was used to identify trans-acting host factors that interact with the 5′-UTR of SARS-CoV-2. Several proteins showed 100% relative binding scores, including A2BP1, ZRANB2, sap-49, FUS, PUM2, SRSF9, MBNL1, KHSRP, RBMX, SFRS1, and HuR (Fig S1A, B). Bioinformatics-based predictions of RBPs with the RBPDB were based on matching linear sequence motifs.

To study the role of HuR in the viral life cycle, ACE-2-expressing HEK-293T cells were transfected with HuR short-interfering RNA (siRNA), which partially knocked down HuR expression by 70% (si HuR) or control siRNA (Nsp si), and infected with SARS-CoV-2 at a multiplicity of infection (MOI) of 0.1. The cells were harvested 48 h post-infection, and the viral titers in the supernatants of siHuR-treated cells and non-specific siRNA-treated cells were compared. We observed that siHuR treatment decreased the viral titer by more than 1 log (Fig 1A, B), emphasizing its important role in the viral life cycle. Parallel experiments were performed to verify the effect of HuR on the life cycle of different viral variants. Silencing of HuR decreased the viral titers of all variants, but the effect was delayed with the alpha variant, with no difference observed after 24 h and a clear difference observed after 48 h (Fig 1C-E). To confirm the role of HuR, we generated HuR-knock out (KO) HEK-293T-ACE-2 cells using a guide RNA targeting the HuR coding region. HuR KO was confirmed by western blotting using an anti-HuR antibody (Fig 1F) and by sequencing (Fig 1G). HuR KO increased the cell proliferation slightly (Fig S2A). We infected wild-type (WT) HuR-positive cells and HuR-KO cells with SARS-CoV-2 at an MOI of 0.1 and examined differences in the supernatant viral titers and cellular SARS-CoV-2 RNA levels. Knocking out HuR reduced the cellular viral RNA levels by >80% (Fig 1H) and decreased the viral titer by >2 logs (Fig 1I). The abundances of SARS-CoV-2 sub-genomic RNAs (sgRNAs) encoding the Spike and Nucleocapsid proteins decreased similar to that of genomic RNA in the HuR KO cell line (Fig 1J). The HuR KO cell line was also utilized to assess the impact of HuR complementation on viral RNA levels in SARS-CoV-2 variants of concerns (VoCs). Overexpression of HuR was found to increase the viral RNA in all the VoCs suggesting an important role in viral life cycle (Fig S2B).

**Fig 1.**
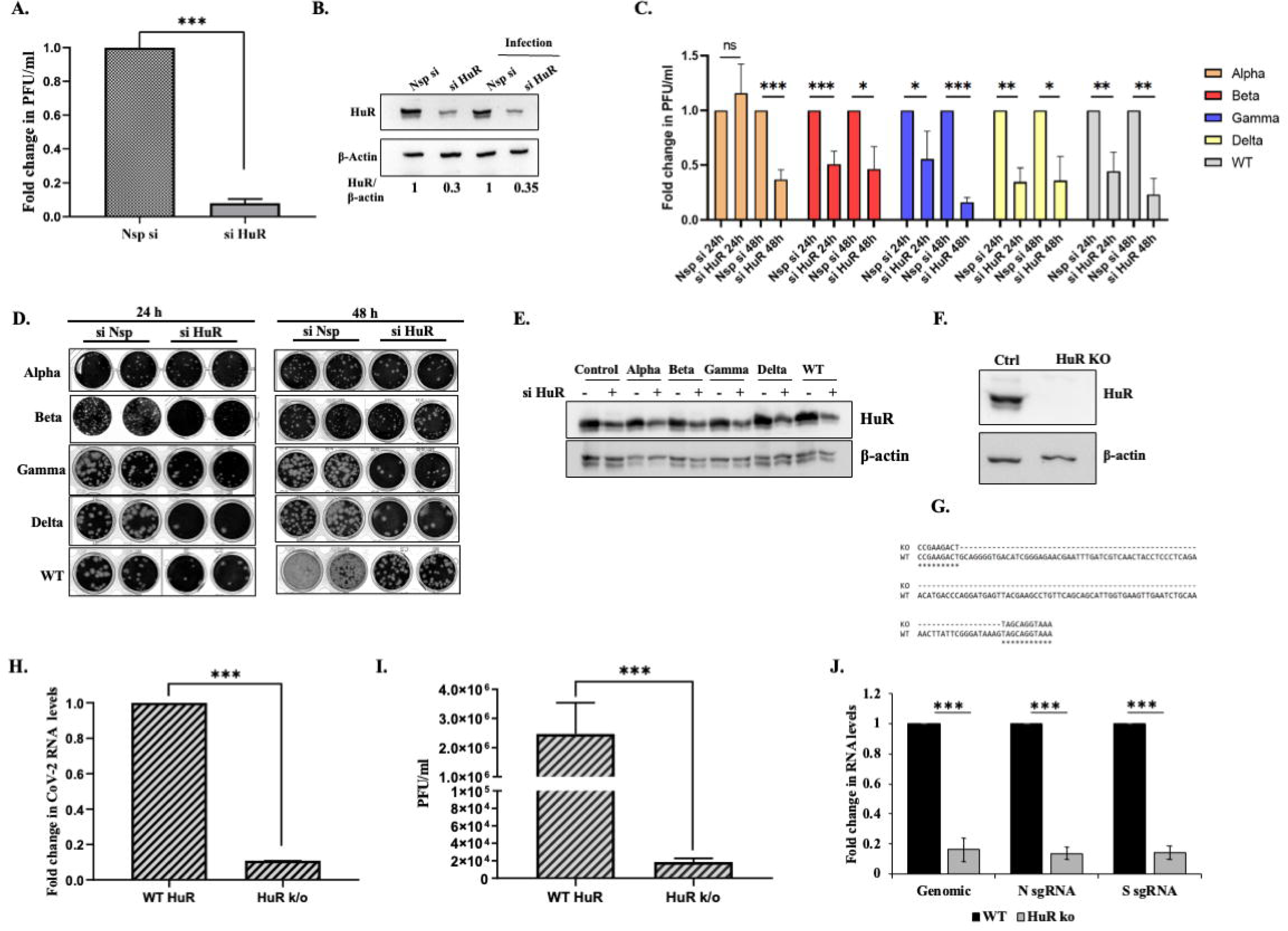
Role of HuR in SARS-CoV-2 lifecycle. (A) HEK-293T-ACE2 cells were transfected with either non-specific (Nsp si) or siRNA targeting HuR. 16 hrs post transfection, cells were infected with 0.1 MOI of SARS-CoV-2 virus. Viral titre in supernatant was determined using plaque assay (n=3). (B) Western blotting was done to confirm HuR silencing using anti-HuR antibody. (C) Effect of HuR silencing on different SARS-CoV-2 VoCs. HEK-293T-ACE2 cells were transfected with either non-specific siRNA (Nsp si) or siRNA targeting HuR (si HuR). 16 hrs post transfection cells were infected with different SARS-CoV-2 VoCs at 0.1 MOI. 48 hrs post infection cell supernatant was collected to determine the viral titres by plaque assay (n=3). (D) Representative images for plaque assay with VOCs. (E) Western blotting using anti-HuR antibody was performed for confirmation of HuR silencing. (F) HuR KO was confirmed by western blotting using anti-HuR antibody. (G) Sequence alignment of HuR gene being targeted by the CRISPR gRNA. The blanks in KO sequence represent the deletion of the region between the two gRNAs. (H) WT HuR and HuR KO HEK-293T-ACE-2 cells were infected with SARS-CoV-2 virus at 0.1 MOI and 48 hrs post infection, positive strand RNA levels were determined using primers specific for SARS-CoV-2 5’UTR from WT and HuR KO cells harvested after 48 hrs of infection (n=3). (I) WT HuR and HuR KO HEK-293T-ACE-2 cells were infected with SARS-CoV-2 virus at 0.1 MOI and 48 hrs post infection, viral titre in supernatant was determined using plaque assay (n=3). (J) RNA levels of genomic 5’UTR and that of sub-genomic RNAs for Spike and Nucleocapsid were quantified using specific primers (n=3). Student t-test was used for statistical analysis. *=p<0.05, **=p<0.01, ***=p<0.001.

### HuR bound the 5′-UTR of SARS-CoV-2

To verify that HuR can bind the SARS-CoV-2 5′-UTR, we performed ultraviolet (UV)-crosslinking assays with a SARS-CoV-2 5′-UTR probe (transcribed *in vitro* with P^32^) and recombinant HuR protein. Direct UV crosslinking showed that HuR bound the labeled SARS-CoV-2 5′-UTR probe in a dose-dependent manner (Fig 2A). The interaction was further confirmed by performing competitive UV-crosslinking experiments, where 100-fold and 150-fold excesses of unlabeled self-RNA showed increasing competition, which was not observed with an unlabeled non-specific RNA (Fig 2B). These results confirmed the *in vitro* binding of HuR to the SARS-CoV-2 5′-UTR.

**Fig 2.**
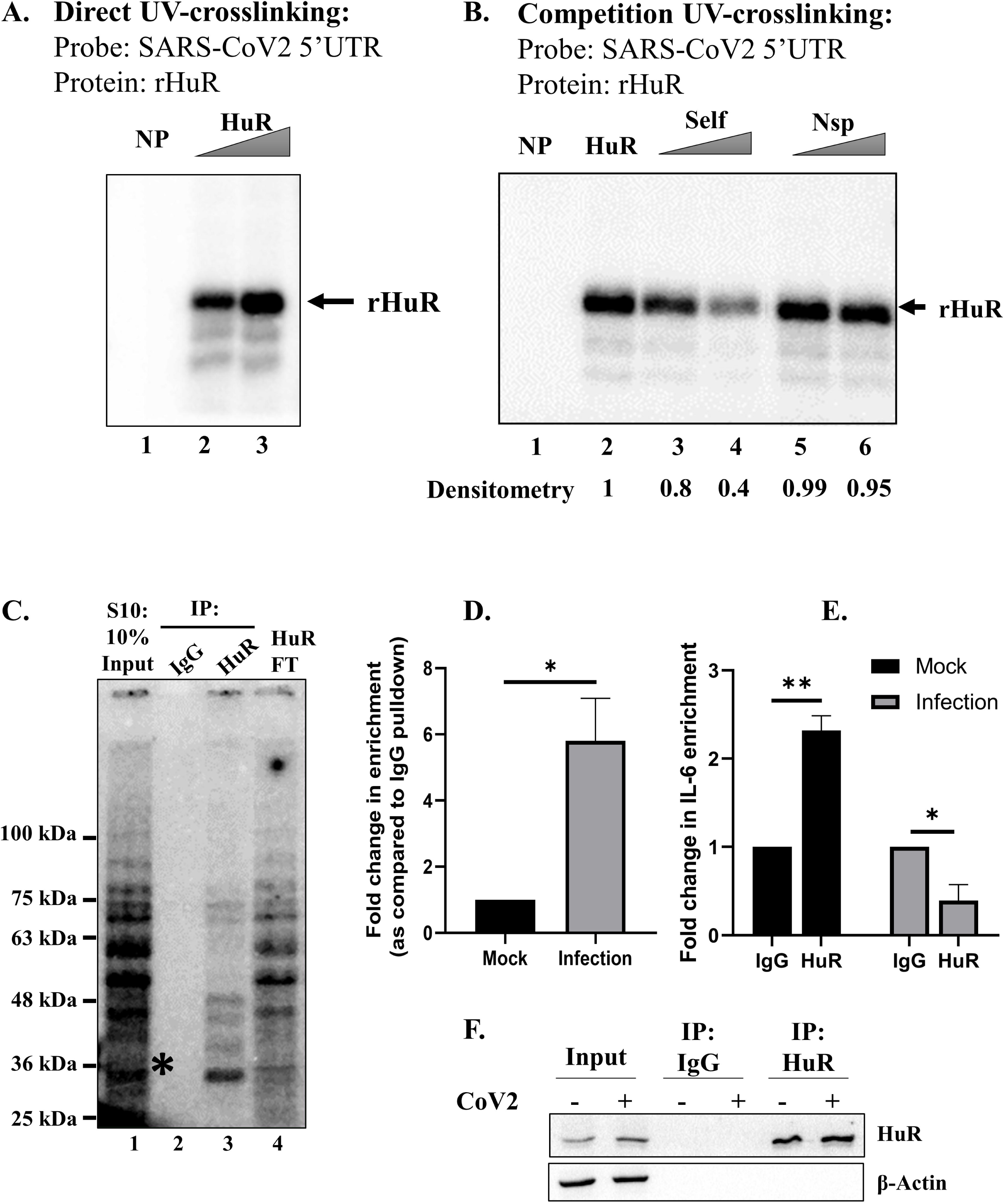
HuR binds to SARS-CoV-2 5’UTR. (A, B) *in vitro* interaction of recombinant HuR with radiolabelled SARS-CoV-2 5’UTR using UV-crosslinking assay. (C) UV-crosslinking of S10 extract with radiolabelled SARS-CoV-2 5’UTR, followed by immune pull down of HuR. The asterisk indicates the band at predicted molecular weight of HuR. (D) *ex vivo* interaction of HuR with CoV-2 5’UTR in SARS-CoV-2 virus infected cells was quantified upon immune-pulldown using anti-HuR antibody. IgG pulldown was used as negative control to calculate the enrichment of SARS-CoV-2 5’UTR in HuR pulldown. Fold change of enrichment was plotted as compared to the enrichment in mock (n=3). (E) *ex vivo* interaction of HuR with IL-6 RNA in mock and SARS-CoV-2 virus infected cells was quantified upon immune-pulldown using anti-HuR antibody. IgG pulldown was used as negative control and the fold enrichment of IL-6 in HuR pulldown was calculated as normalised to IgG pull down (n=3). (F) The western blot for HuR pull-down in panel D and E. Student t-test was used for statistical analysis. *=p<0.05, **=p<0.01, ***=p<0.001.

To verify the binding in cell lysates, we performed UV-crosslinking immunoprecipitation (IP) experiments wherein HuR was pulled down from the cytoplasmic S10 lysates following UV crosslinking with the SARS-CoV-2 5′-UTR, using an anti-HuR antibody. Specific HuR binding to the SARS-CoV-2 5′-UTR was demonstrated by comparing the electrophoretic bands between experimental (after crosslinking and IP) and input samples. The band at approximately 36 kDa, which was absent from the flow-through beads, depicted the HuR band in the cell lysate (Fig 2C). *Ex vivo* binding of HuR to the SARS-CoV-2 5′-UTR in infected cells was demonstrated by performing IP-reverse transcription (RT) experiments where HuR was pulled down using an anti-HuR antibody and the associated SARS-CoV-2 5′-UTR was quantified using specific primers. The enrichment of SARS-CoV-2 5’UTR in the HuR pull down upon infection confirms the *ex vivo* interaction (Fig 2D). IgG pull down was used as a negative control for HuR pull down. IL-6 RNA, which is known interactor of HuR [18] was used as a positive control. We observed significant enrichment of IL-6 in HuR pull down fraction from mock cells (Fig 2E). This interaction reduced upon SARS-CoV-2 infection, which could be because of HuR sequestration by the viral RNA.

### HuR bound around the C241 site in the SARS-CoV-2 5′-UTR

To identify the HuR-binding site in the SARS-CoV-2 5′-UTR, we used a catRAPID fragment software that considers the secondary structure when determining the binding affinity of a protein. HuR showed very high affinity for the region spanning nts 150-250, which corresponds to stem-loop 5 of the SARS-CoV-2 5′-UTR (Fig S3). Two HuR-binding motifs (199-GUUU-202 and 237-GUUU-240) were found within the region spanning nts 150-250 nts, which overlapped with the loop regions of SL5A and SL5B, respectively. However, the catRAPID fragment showed lower affinity for the third 8-GUUU-11 motif, which base paired with the SL1 hairpin.

To assess the impact of HuR binding, we evaluated the presence of mutations in the 5′-UTRs of different SARS-CoV-2 variants. We aligned the 5′-UTRs of the alpha, beta, gamma, and delta variants and found two mutations between them. The alpha variant acquired a C241U mutation, which was retained in the other successive variants. The beta and delta variants acquired another mutation, namely G210U (Fig 3A).

**Fig 3.**
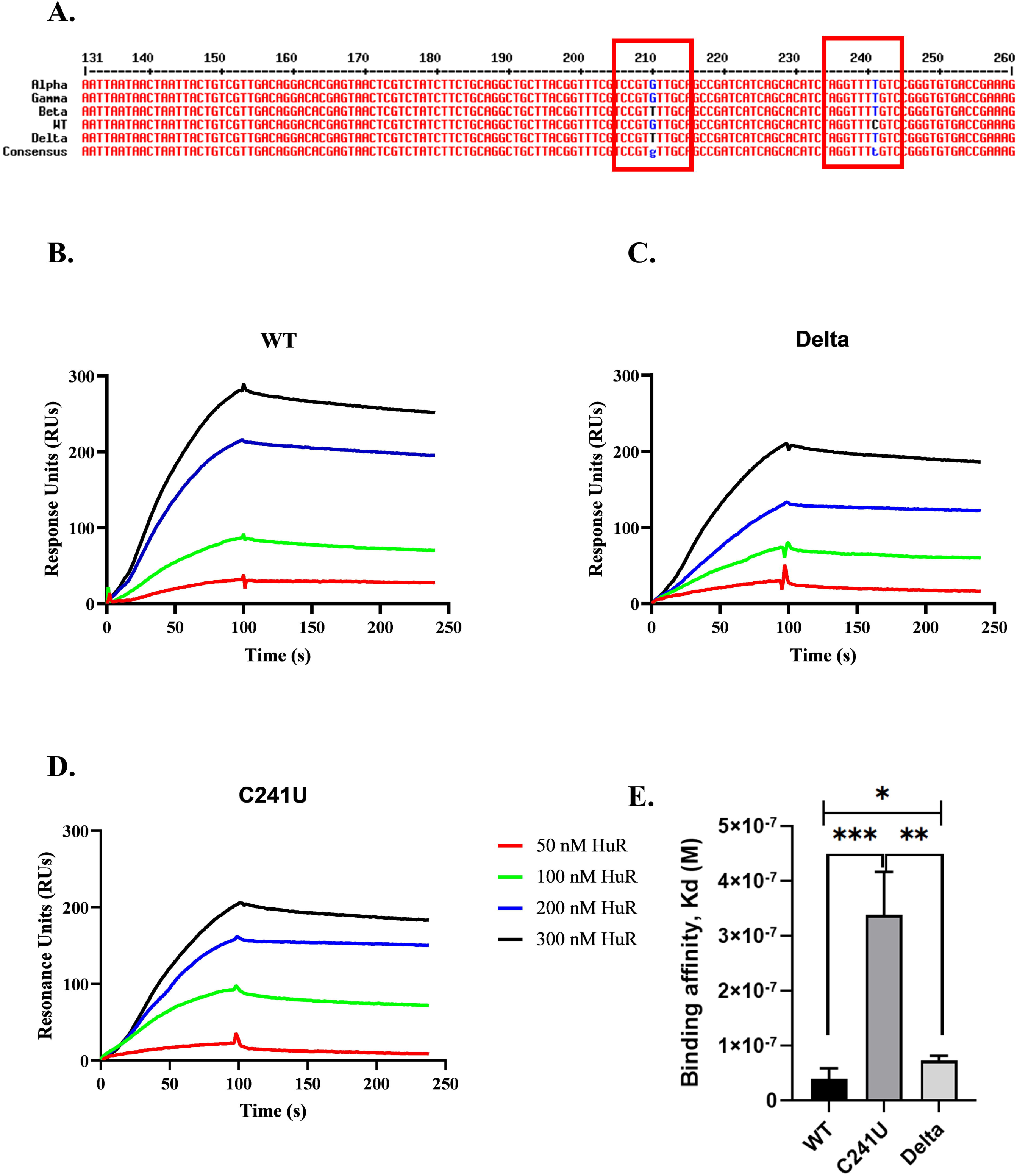
HuR binding site on SARS-CoV-2 5’UTR. (A) Multiple sequence alignment of 5’UTR of alpha, beta, gamma and delta variants of SARS-CoV-2. (B, C, D) Sensorgrams for indicated biotinylated SARS-CoV-2 5’UTR binding to recombinant HuR using SPR. Colors denote increasing HuR concentration (E) Binding affinity of mutants obtained by SPR. (n=3). Student t-test was used for statistical analysis. *=p<0.05, **=p<0.01, ***=p<0.001.

We performed Surface Plasmon Resonance (SPR) studies to analyze the effect of these mutations on the HuR-binding affinities (Fig 3B-D). Incorporation of the C241U mutation decreased the binding affinity of HuR from 32.5 +/- 13 nM (WT strain) to 338.6 +/- 63 nM, whereas the incorporation of G210U along with C241U increased back the binding affinity to 72.9 +/- 7.2 nM (Fig 3E). These results could indicate that the second mutation (G210U) generated an additional binding site for HuR. These findings corroborate the variant data shown in Figure 1, and the lower binding affinity of HuR to the alpha variant might explain its delayed effect. Furthermore, the 5′-UTRs of the alpha and gamma variants contained the same mutations, but the gamma variant did not show lesser HuR dependency at 24h post infection. The results could reflect other direct or indirect effects of HuR silencing. These data provide insights into the differential roles of HuR in SARS-CoV-2 variants due to the altered binding affinity of HuR. Furthermore, we analysed the sequence conservation of these HuR binding sites at 210 and 241 position amongst the beta coronaviruses, MERS, SARS-CoV and SARS-CoV-2. The 5’UTRs of SARS-CoV and SARS-CoV-2 exhibited high sequence conservation (Fig S3C). We observed that the HuR binding site around 241 position is completely conserved in the beta coronaviruses and the binding site around 210 position showed a mutation from “C” in MERS to “G210” in SARS-CoV and SARS-CoV-2 (Fig S3C). The C to G mutation might not alter the HuR binding drastically at the described site. The evolutionary conservation of HuR binding site in the 5’UTR highlights its importance in viral life cycle and propagation.

### HuR promoted translation from the SARS-CoV-2 5′-UTR

SARS-CoV-2 replication can be impacted through the regulation of many different steps in the viral life cycle. Because HuR bound to the SARS-CoV-2 5′-UTR (where translation of the genome begins), we investigated whether HuR regulates this step. To this end, we generated a luciferase reporter construct wherein the viral 5′-UTR was followed by the luciferase gene. Luciferase activities under different conditions revealed various degrees of SARS-CoV-2 5′-UTR translation.

A549 cells were transfected with a 5′-UTR-luc construct in the context of either partial HuR knockdown (Fig 4A) or HuR overexpression (Fig 4B), and the luciferase activities were measured. To normalize the luciferase activities, the cells in each well were co-transfected with 2 ng of pRLTK (encoding Renilla luciferase), and dual-luciferase readings were taken after 8 or 12 h. Partial HuR knockdown reduced translation from the SARS-CoV-2 5′-UTR (Fig 4A), and HuR overexpression increased translation (Fig 4B). These results suggest that HuR promoted viral RNA translation by binding to the 5′-UTR. This effect was also observed in HuR-KO cells. The inclusion of SARS-CoV-2 5′-UTR in luciferase reporter construct showed increased luciferase activity in WT cells (lanes 1 and 2), suggesting that the presence of the SARS-CoV-2 5′-UTR might have recruited additional factors needed for enhancing cap-dependent viral translation. Furthermore, this increase was completely abrogated in HuR KO cells (lanes 3 and 4), implying the importance of HuR as the factor behind increased translation from the viral 5′-UTR. We quantified the luciferase RNA levels in this experiment and observed no significant difference, suggesting the impact of HuR on viral RNA translation (Fig S4A). Notably, translation from the parental luciferase construct was reduced in HuR KO cells, which could represent an independent effect of HuR knockdown (Fig 4C).

**Fig 4.**
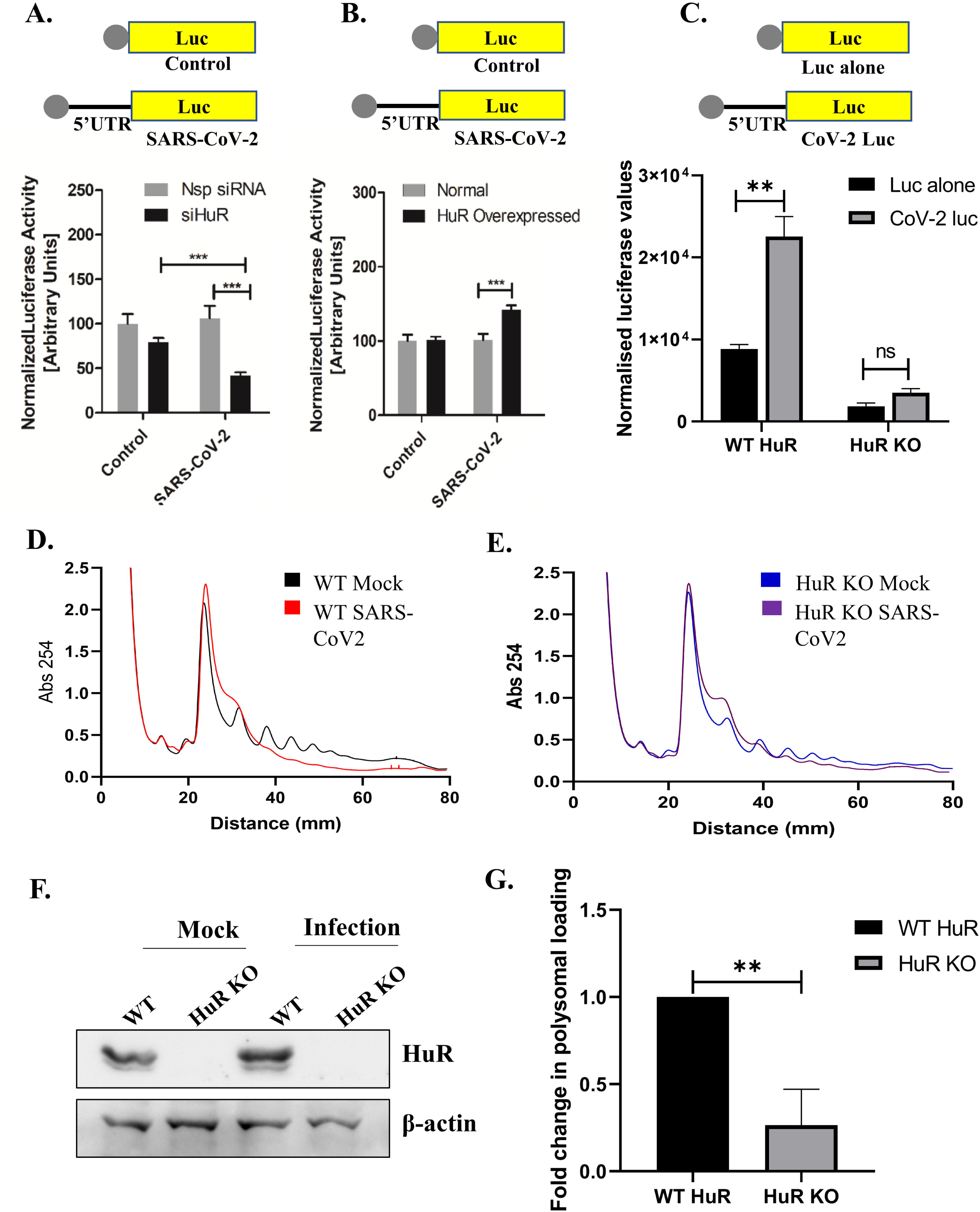
HuR promotes SARS-CoV-2 translation by binding to its 5’UTR. (A) A549 cells were transfected with either non-specific (Nsp si) or siRNA targeting HuR (siHuR). 16 hrs post transfection, cells were transfected with either pGL3-SV40-Luc (Control) or pGL3-SV40-SARS-CoV-2 5’UTR-Luc (SARS-CoV-2 Luc), depicted as schematics above the graphs, and harvested after 12 hrs of transfection and luciferase readings measured. pRLTK was co-transfected for transfection control and used for normalization of luciferase values. Effect of si HuR was calculated with respect to the luciferase in Nsp si condition (n=3). (B) HuR overexpression was performed in place of silencing in the previous panel. Effect of overexpression was calculated with respect to luciferase in no overexpression condition (n=3). (C) WT HuR and HuR KO HEK-293T-ACE2 cells were transfected with either pGL3-SV40-Luc (Luc alone) or pGL3-SV40-SARS-CoV-2 5’UTR-Luc (CoV-2 Luc), depicted as schematics above the graphs and luciferase values were measured 24 hrs post-transfection. The graph represents absolute values obtained upon normalization of luciferase readings with protein concentration (n=3). (D) WT HEK-293T-ACE2 cells were infected with SARS-CoV-2 virus at 0.1 MOI and 48 hrs post infection cells were harvested for sucrose gradient polysome fractionation. The lines represent the absorbance at 254 nm v/s the distance from the top of gradient. (E) HuR KO HEK-293T-ACE2 cells were infected with SARS-CoV-2 virus at 0.1 MOI and 48 hrs post infection cells were harvested for sucrose gradient polysome fractionation. The lines represent the absorbance at 254 nm v/s the distance from the top of gradient. (F) Western blotting was performed for the lysates used for polysome fractionation using anti-HuR antibody. (G) RNA was isolated from monosome and polysomal fractions (separately) of infected lysate from panel (D) and (E) and the abundance of SARS-CoV-2 5’UTR quantified by specific primers in real time PCR. Polysomal loading was calculated as the ratio of RNA in Polysome:Monosomes (n=3). Student t-test was used for statistical analysis. *=p<0.05, **=p<0.01, ***=p<0.001.

The role of HuR in SARS-CoV-2 translation was further studied by analyzing viral RNA loaded into the polysomes in WT and HuR-KO cells. The reduction in polysomal peaks upon SARS-CoV-2 infection suggested that SARS-CoV-2 infection led to a global reduction in translation (Fig 4D). This reduction was compromized in HuR-KO cells (Fig 4E). We compared the global translation in WT and HuR KO cells by calculating the ratio of the area of polysomal to monosomal peak. Higher Polysome: Monosome ratio suggests higher translation. In all three experimental sets, we observed a slightly reduced Polysome: Monosome ratio in HuR KO cell line as compared to the WT, but the difference was not statistically significant (Fig S4B). This suggests that there might be slight reduction in global translation in HuR KO cell, which might also explain the lower luciferase value of the Luc alone vector control in the HuR KO cell as compared to WT cells. We also calculated the Polysome: Monosome ratio in infected WT and HuR KO cells, where we found higher ratio in the HuR KO cells (Fig S3C). This further confirms that the global translational reduction upon SARS-CoV-2 infection is much higher in WT cells than in HuR KO cells. Of note, we cannot compare the ratio of area under the curve (AUC) in Mock and infected cells because of the major differences in the polysome profiles in these conditions such as the transition phase from monosome to polysomes (Fig 4D, E). Further, to assess the polysomal loading of the viral RNA, RNA was isolated from monosome and polysome fractions separately, and the abundances of the SARS-CoV-2 5′-UTR were quantified by real-time polymerase chain reaction (PCR) analysis using specific primers. The RNA loading to polysomes was calculated as a ratio of RNA in Polysome: Monosomes. We found increased polysomal loading of SARS-CoV-2 RNA as compared to the actin RNA in WT infected cells, suggesting the preferential loading of viral RNAs in the polysome even after global translational reduction (Fig S4D). HuR knockout led to a 60-70% decrease in SARS-CoV-2 RNA loading into polysomes as calculated by ratio of RNA loaded in polysome: monosomes (Fig 4F, G). This finding confirms the important role of HuR in promoting translation from SARS-CoV-2 5′-UTR.

### HuR regulated the translation of SARS-CoV-2 sub-genomic RNAs

We observed that HuR KO led to decreased viral genomic and sgRNA production (Fig 1J). SARS-CoV-2 possesses 2 viral RNA species during its life cycle [19]. One is the genomic RNA which has a 272 nts long 5’UTR upstream to the ORF coding for non-structural proteins. After viral replication from 3’end, the second species of RNA, known as sub-genomic RNAs (sgRNAs) are generated. These code for individual viral structural and accessory proteins. The 5’UTR of sgRNAs possess only 1-69 nts of the genomic 5’UTR [20]. Since they are independent RNA species with only first 69 nts overlap in the 5′-UTR[21], we examined the effect of HuR KO on translation on the sg 5′-UTR. To this end, we infected WT and HuR KO cells with SARS-CoV-2 at an MOI of 1.0 and harvested the cells and culture supernatants after 24 h. We observed a >10-fold reduction in viral titers in HuR-KO cells (Fig 5A). Furthermore, the genomic and sub-genomic RNA levels decreased to the same extent in HuR-KO cells (Fig 5B). We quantified the expression levels of non-structural and structural proteins (produced by the translation of genomic and sub-genomic 5′-UTRs, respectively) in WT and HuR-KO cells by western blot analysis. The Nucleocapsid protein represents sub-genomic translation, and the Nsp5 protein represents genomic translation. We observed that the Nucleocapsid: Nsp5 protein ratio was higher in HuR-KO cells than in WT cells (Fig 5C), even though no difference was evident in their RNA levels. These findings suggested that both RNA species were differentially regulated at the level of translation. We also found a similar effect of HuR KO on another non-structural protein, Nsp1 (Fig S5A), confirming the role of HuR in mediating polyprotein translational from 5’UTR and ruling out the effect on stability of independent non-structural proteins. To study sub-genomic RNA translation, we performed capped *in vitro* transcription of sub-genomic SARS-CoV-2 5′-UTR (nts 1-70) followed by luciferase RNA and used *in vitro*-transcribed luciferase RNA alone as a control. These RNAs were transfected into WT and HuR-KO HEK-293T-ACE-2 cells, and luciferase activities were measured at 8 h post-transfection. As observed previously, transfecting the parental luciferase reporter RNA alone (Luc alone) exhibited lower readings in the HuR-KO cell line. This decrease could be because of the global translational reduction in HuR KO cell line and should have been observed for sg5’UTR luc RNA as well if HuR had no specific role in its translation. However, this decrease was rescued for sg 5′-UTR-luc, demonstrating that the absence of HuR in the KO cell line supported translation through sg5’UTR (Fig 5D). Therefore, we conclude that HuR suppresses the translation of SARS-CoV-2 sub-genomic RNAs, contrary to its role in promoting the translation of SARS-CoV-2 genomic RNAs (Fig 4). We determined the luciferase RNA levels in this experiment and observed no significant difference, suggesting the impact of HuR on viral sgRNA translation (Fig S5B). These data explain the increased ratio of structural to non-structural proteins in the HuR-KO cell line and provide the first evidence of differential translational regulation of SARS-CoV-2 genomic RNA and sgRNA by a host protein.

**Fig 5.**
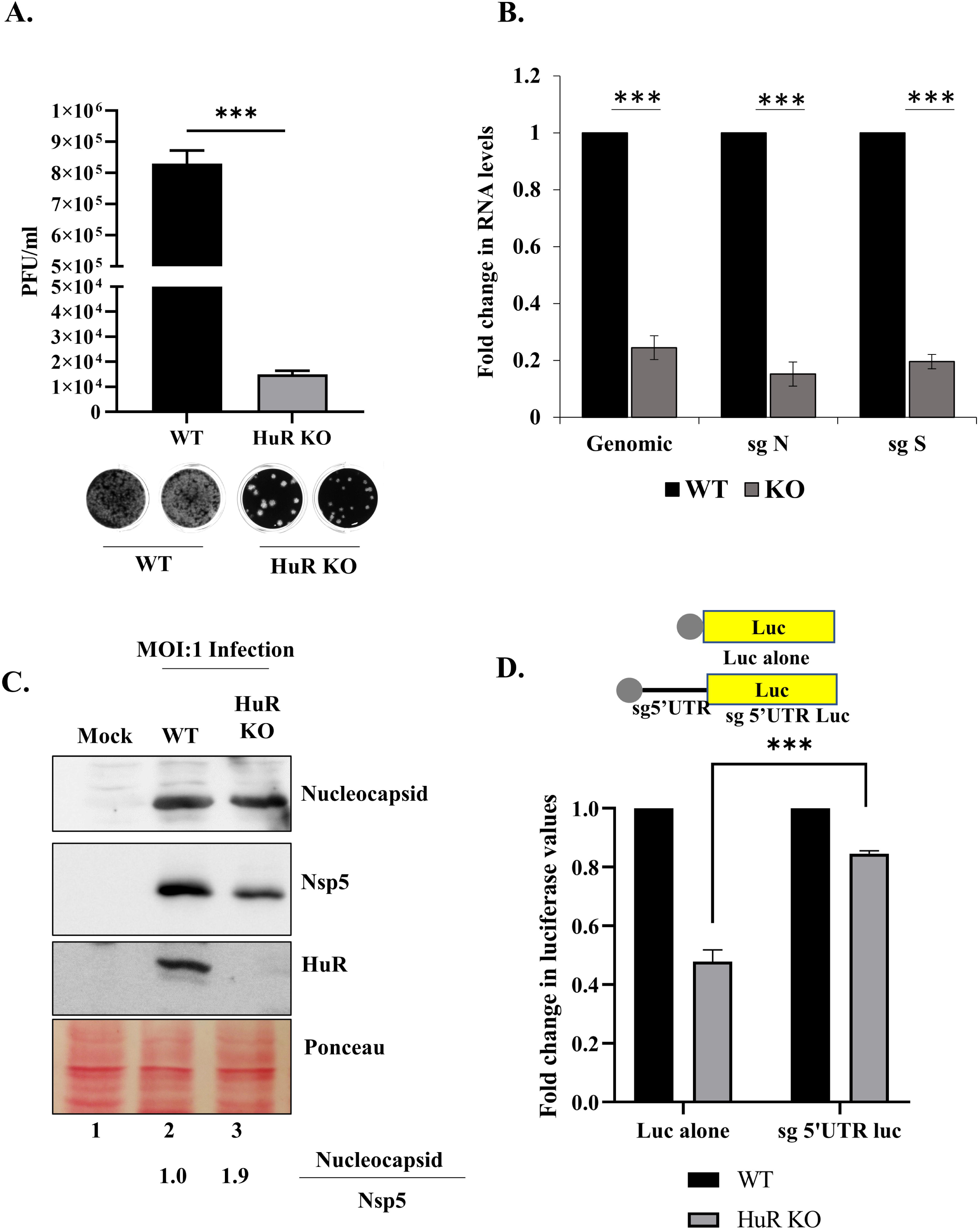
HuR inhibits translation of SARS-CoV-2 sub-genomic RNAs. (A) WT HuR and HuR KO HEK-293T-ACE2 cells were infected with SARS-CoV-2 virus at MOI of 1.0 and 24 h post infection, viral titre in supernatant was determined using plaque assay (n=3). The below panel shows representative plaque assay images for the same. (B) RNA levels of genomic 5’UTR and that of sub-genomic RNAs for Spike and Nucleocapsid were quantified in the same experiment using specific primers (n=3). (C) Protein levels of Nucleocapsid and Nsp5 were determined in the above experiment by western blotting using specific antibodies. Ponceau staining of the bot represents equal loading. The values at the bottom represent the average densitometry values after quantifying the intensities of Nucleocapsid and Nsp5 bands (n=3). (D) WT HuR and HuR KO HEK-293T-ACE2 cells were transfected with either *in vitro* transcribed capped Luciferase RNA (Luc alone) or *in vitro* transcribed capped sub-genomic 5’UTR followed by Luciferase RNA (sg 5’UTR luc), as depicted in schematic above the graph, and luciferase values were measured 8 h post-transfection. The graph represents Fold change in luciferase values obtained upon normalization of luciferase readings with protein concentration (n=3). Student t-test was used for statistical analysis. *=p<0.05, **=p<0.01, ***=p<0.001.

### HuR regulated SARS-CoV-2 translation by assisting binding of the polypyrimidine tract-binding (PTB) protein

To further investigate the mechanism of action of HuR, we analyzed the binding of different cellular proteins to the 5′-UTR of SARS-CoV-2. We purified cellular S10 extracts from three different cell lines (HEK-293T-ACE-2, A549, and Huh7.5) and analyzed their binding to the SARS-CoV-2 5′-UTR by performing UV-crosslinking experiments. We observed that multiple proteins bound to the 5′-UTR in a dose-dependent manner (Fig 6A). The band marked with an asterisk has a mass that corresponded to the size of HuR, as verified in UV-IP experiments (Fig 2C).

**Fig 6.**
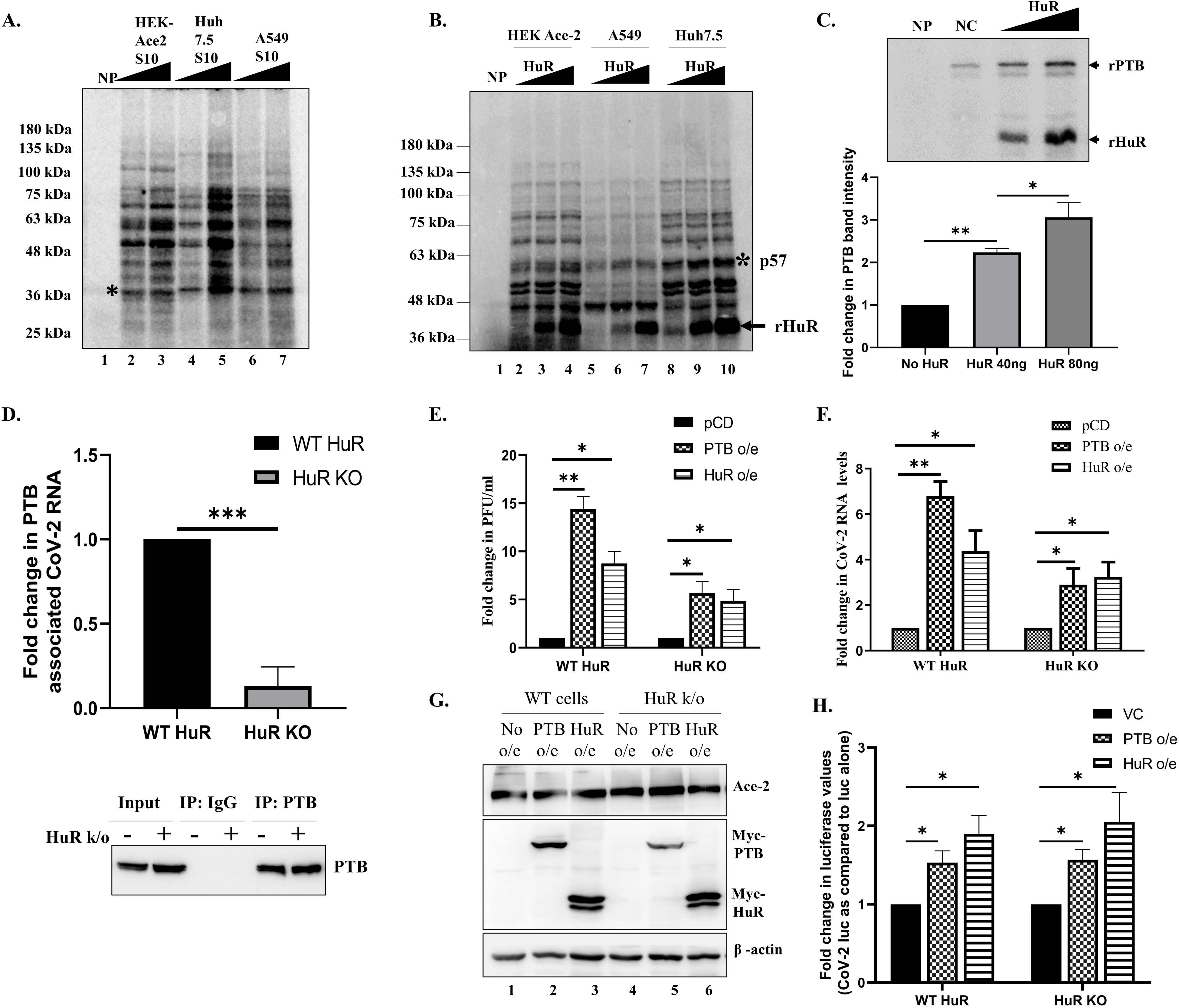
HuR assists PTB binding to 5’UTR to promote translation. (A) UV-crosslinking profiles with increasing amount of S10 extracts of different cell lines as indicated. Asterisk indicates the band at the size of HuR. (B) UV-crosslinking was performed with S10 extracts of different cell lines and increasing amount of HuR. Asterisk indicates the band at the size of PTB. (C) Competition UV-crosslinking was performed with constant amount of PTB and increasing amounts of recombinant HuR. The lower panel represents the densitometry of PTB band intensity (n=3). (D) HEK-293T-ACE-2 with WT and HuR KO were infected with 0.1 MOI of SARS-CoV-2. 48 hrs post infection, cells were harvested and PTB immunoprecipitated. The associated RNA was quantified by real time PCR using SARS-CoV-2 5’UTR specific primers (Upper panel). IgG pulldown was used as negative control to calculate the enrichment of SARS-CoV-2 5’UTR in PTB pulldown. Fold change of enrichment was plotted as compared to the enrichment in WT cells (n=3). The immune pull down was confirmed by western blot using anti-PTB antibody (Lower panel). (E) WT and HuR KO HEK-293T-ACE2 cells were transfected with myc-tagged HuR and PTB overexpression constructs. 16 hrs post-transfection, cells were infected with SARS-CoV-2 at 0.1 MOI. Cell supernatant was collected after 48 hrs of infection and Viral titres determined using plaque assay (n=3). (F) Viral RNA levels were quantified in cells from the previous experiment (n=3). (G) Western blotting was performed using anti-myc antibody confirmation of overexpression. (H) WT and HuR KO HEK-293T-ACE2 cells were transfected with myc-tagged HuR and PTB overexpression constructs. 16 hrs post-transfection, cells were transfected with pGL3-SV40-Luc (Luc alone) or pGL3-SV40-SARS-CoV-2 5’UTR-Luc (CoV-2 Luc) and harvested after 24 hrs of transfection and luciferase readings measured (n=3). Student t-test was used for statistical analysis. *=p<0.05, **=p<0.01, ***=p<0.001.

We checked whether the addition of exogenous HuR to the S10 lysate would alter the binding to any other protein and observed a band, approximately 57 kDa in size, whose intensity increased with increasing concentrations of HuR (Fig 6B). We previously characterized PTB as an important protein with a band in this size range[9]. Therefore, we purified recombinant PTB and HuR proteins and performed competitive-crosslinking experiments with increasing HuR concentrations, while keeping the PTB concentration constant. HuR binding promoted PTB binding to the SARS-CoV-2 5′-UTR in a concentration dependent manner, corroborating our previous data (Fig 6C). We analyzed whether this competition occurs *ex vivo* in cells by performing IP-RT experiments, where PTB was immunoprecipitated from cells expressing WT HuR and HuR-KO cells (Fig 6D). Thereafter, the associated 5′-UTR was quantified by real time PCR. The reduction in PTB associated viral RNA in the absence of HuR (Fig 6D, upper panel), confirms that HuR functions by promoting the binding the PTB.

Because HuR is primarily a nuclear protein that relocalizes to the cytoplasm upon infection with different viruses, we checked the localization of HuR at different time points following infection by conducting immunofluorescence and nuclear-cytoplasmic fractionation experiments (Fig S6). In the immunofluorescence assays, HuR localized to the nucleus in mock-infected cells, and we did not observe any changes in HuR localization when HEK-293T-ACE-2 cells were infected with SARS-CoV-2 at different timepoints (Fig S6A). Nuclear-cytoplasmic fractionation experiments revealed that a fraction of HuR localized to the cytoplasm and that its cytoplasmic abundance did not increase upon SARS-CoV-2 infection. These findings suggest that cytoplasmic HuR is sufficient for promoting PTB binding and influencing viral translation.

We then examined whether PTB overexpression could rescue the decrease in viral titer in HuR-KO cells. PTB overexpression increased the viral titer by more than 1 log in both the WT HuR and HuR-KO cell lines (Fig 6E). PTB overexpression demonstrated higher increase in viral RNA levels and viral titres in WT HuR cell line as compared to the HuR KO cell line suggesting that the effect of PTB on viral life cycle is mediated through HuR (Fig 6F, G). The increase in viral RNA and titres upon PTB overexpression was slightly higher than HuR overexpression which could be because of the inhibitory effect of the HuR on sgRNAs. Luciferase constructs were used to examine whether these effects were mediated through translational regulation. PTB overexpression increased luciferase activity in WT and HuR KO cell line (Fig 6H). Slightly higher increase in translation was observed upon HuR overexpression. These results suggest that the presence and binding of HuR to the SARS-CoV-2 5′-UTR could guide PTB binding and thereby promote viral translation. The same effect was observed by increasing PTB expression in cells, which would increase its binding propensity and affinity because of the increased abundance.

### HuR can potentially be used as a target for developing better therapeutics

Because HuR played an important role in guiding SARS-CoV-2 viral translation by binding to viral RNA, we explored whether it could alter the activity of remdesivir in viral inhibition. We used a concentration gradient of remdesivir and examined the decrease in viral RNA in WT and HuR-KO cells. We found that 200 nM remdesivir completely inhibited viral RNA production in both cell lines; however, at lower remdesivir concentrations, the HuR-KO cell line showed substantially lower viral RNA levels (Fig 7A). The half-maximal inhibitory concentration (IC_50_) of remdesivir was calculated based on the resulting data, and we found that the IC_50_ was 10.46 nM in WT cells but only 1.82 nM in HuR-KO cells (Fig 7B). These data suggest the potential of targeting HuR as an additional therapy in combination with remdesivir.

**Fig 7.**
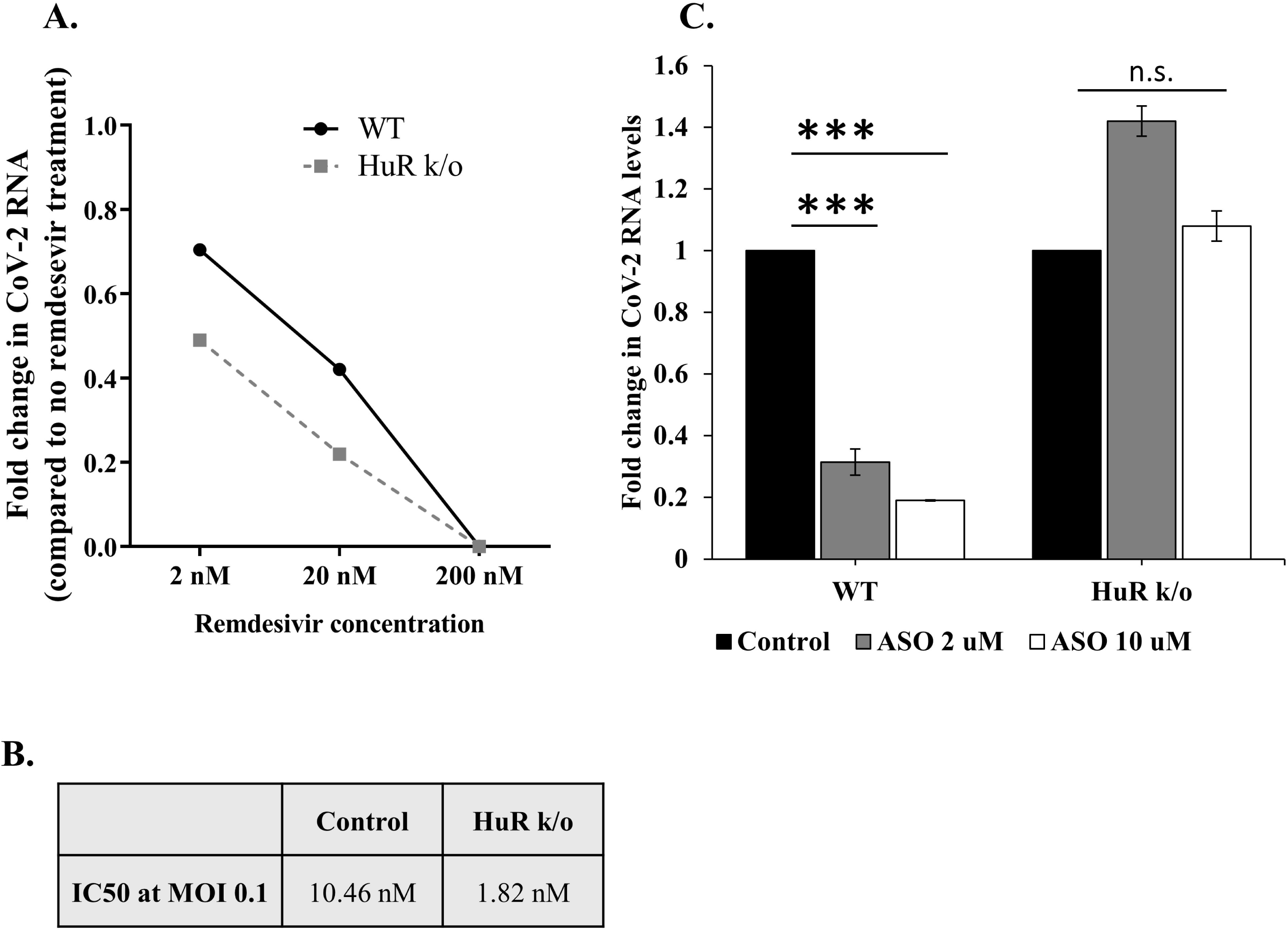
Therapeutic significance of the role of HuR in SARS-CoV-2 life cycle. (A) Effect of HuR KO on remdesivir sensitivity of viral infection. WT and HuR KO HEK-293T-ACE-2 cells were infected with SARS-CoV-2 virus at an MOI of 0.1 and the mentioned concentrations of Remdesivir added after 1 hr of virus inoculation. Cells were harvested after 48 hrs of treatment and RNA isolated to detected SARS-CoV-2 5’UTR. (B) Table for IC50 of remdesivir for CoV-2 infection in WT and HuR KO cell line. (C) Effect of anti-sense oligos (ASOs) targeting HuR binding site on viral titres. WT and HuR KO HEK-293T-ACE-2 cells were transfected with the mentioned concentrations of ASO and 12 hrs post transfection, cells were infected with SARS-CoV-2 virus at 0.1 MOI. 48 hrs post infection, cells were harvested, and SARS-CoV-2 RNA quantified using real time PCR (n=3). Student t-test was used for statistical analysis. *=p<0.05, **=p<0.01, ***=p<0.001.

Recently, ASOs directed against different regions of SARS-CoV-2 RNA have been demonstrated to inhibit SARS-CoV-2 replication in cell culture and in animal models [22]. Of the ASOs used, ASO5 targeted the HuR-binding site (encompassing 241 site) and reduced SARS-CoV-2 RNA levels. Therefore, we investigated the activity of ASO5 using WT and HuR-KO cell lines. As observed previously, ASO5 transfection reduced viral RNA levels in a dose-dependent manner in WT cells. However, such a decrease was not observed in the HuR-KO cell line (Fig 7C), strengthening the evidence supporting the location of the HuR-binding site, its role in the viral life cycle, and its use as a potential therapeutic target.

## Discussion

Host proteins are important factors that regulate the virus life cycle. RNA-binding proteins bind to viral RNAs at different stages of their life cycles, thereby modulating their cellular interactions. RBP screening has identified common proteins involved in the life cycle of multiple viruses. HuR is one such protein that has been shown to regulate the translation and replication of viruses, including hepatitis C Virus[9], coxsackievirus[11], and Sindbis virus[13]. We studied the role of this important protein in regulating the SARS-CoV-2 life cycle. We found that HuR bound to the viral 5′-UTR and regulated its life cycle by promoting genomic 5′UTR mediated translation. It did so by promoting the binding of another RBP, PTB, which can enhance ribosome recruitment to the viral 5′-UTR. HuR binding occurred near 241 site in the 5′-UTR, which is mutated in some SARS-CoV-2 variants of concern. This mutation alters the binding affinity for HuR and the reduces the dependency of alpha variant on HuR, which could be one of the mechanisms behind the longer generation time and hence lesser transmission of the alpha variant as compared to the delta variant[23]. Knocking out HuR increased the sensitivity of infected cells towards remdesivir-mediated viral inhibition. This suggests the potential of HuR inhibition to be used as an add-on therapy for inhibition. The HuR-binding site lies in the target region of anti-sense oligo ASO5[22], which inhibits the viral life cycle and therefore is an attractive candidate for add-on therapy. We further addressed the HuR-mediated translational regulation of SARS-CoV-2 genomic and sgRNAs. We showed that while HuR promoted translation from the genomic 5′-UTR, it inhibits sg 5′-UTR translation. Previous reports have demonstrated the translational regulation by viral proteins such as Nsp1[24, 25] and ORF6[26], and we propose that HuR could represent the cellular arm of such regulation. The genomic and sub-genomic 5′-UTRs vary in the length. The sub-genomic UTR harbors one of the 4 predicted HuR binding sites (8-10) and that site resides in the stem region of the secondary structure while the other 3 sites, which are present only in the full length UTR reside in the loop region in the secondary structure [16]. This could be responsible for the differential effect of HuR binding. Furthermore, it could also affect the PTB recruitment capability of HuR, altering the translation regulation at the initial stage of viral infection. Both genomic and sub-genomic RNA have been found to bind to HuR (ELAVL1) and PTB (PTBP1) in genome wide interaction screens [27]. In fact, at the late time point of SARS-CoV-2 infection cycle, PTB binding to sub-genomic RNA was found to be higher than the genomic RNA. Combining this scheme with our translational results suggests that the cell could remodel PTB to act as an anti-viral factor at the later stages of viral infection. The reduced genomic RNA binding would decrease genomic translation and enhanced sub genomic binding would decrease sub-genomic translation, leading to an overall decrease inhibitory role on viral life cycle.

HuR does not relocalize to the cytoplasm upon SARS-CoV-2 infection in HEK-293T-ACE-2 cells, but the small amount of HuR present in the cytoplasm due to nuclear-cytoplasmic shuttling might be sufficient to promote PTB binding, which appears to relocalize to the cytoplasm upon infection. This possibility was supported by our competition-crosslinking assays, where a small quantity of HuR greatly enhanced PTB binding. Furthermore, HuR is known to alter the stability of cellular transcripts. The cytokine storm and disease severity observed upon SARS-CoV-2 infection could be regulated by HuR by virtue of its binding to mRNAs encoding pro-inflammatory cytokines such as interleukin (IL)-6 and IL-8[28, 29]. We also observed that HuR levels increased following SARS-CoV-2 infection. The regulatory mechanism could be interesting, because miRNAs, especially miR-125b-5p, have been shown to target HuR translation[30]. Several small molecules and drugs have been designed which abrogate HuR function by inhibiting the binding of HuR to its cellular transcripts[31, 32]. Many of them have shown excellent efficacy in animal model and some are in human trials as well. Suramin is one such FDA-approved drug which is prescribed for the treatment of African sleeping sickness and is effective against viruses like HIV, Chikungunya and Ebola[33]. It has been shown to be efficacious in preventing SARS-CoV-2 infection as well[34, 35]. Such drugs are important targets to be explored for either their individual or synergistic repurposing as antivirals for SARS-CoV-2 infection.

Taken together, our observations provide important insights and open multiple avenues for targeting the SARS-CoV-2 life cycle by modulating an important host factor, HuR. Targeting HuR through ASOs could be an important add-on treatment along with remdesivir. It would prevent HuR binding to the genomic 5′-UTR while increasing its availability to bind to the sub-genomic SARS-CoV-2 5′-UTR leading to decreased structural protein translation and thereby help in clearing SARS-CoV-2 infection.

## Materials and Methods

### Plasmids and constructs

To make the template for *in vitro* transcription, the SARS-CoV-2 5’UTR was amplified with 5’AATT<u>GCTAGC</u>ATTAAAGGTTTATACCTTCCCAGGTA3’ and 5’AATT<u>AAGCTT</u>CTCGTTGA AACCAGGGACAAG3’ from pCBB 5’UTR (Kind gift from Dr. Milan Surjit, THSTI) and subcloned at the downstream of T7 promoter in NheI and HindIII cloning site of pcDNA3.1+. For luciferase assay, SARS-CoV-2 5’UTR was amplified with 5’AATT<u>AAGCTT</u>ATTAAAGGTTTATACCTTCCCAGGTAAC3’ and 5’AATT<u>CCATGG</u>TTCTCGTTGAAACCAGGGACAAG 3’ to subclone in pGL3 control vector.

pcDNA3.1-myc-HuR and pET28a-HuR were generated as described previously.

pcDNA3.1-myc-PTB was a kind gift from Dr. Douglas Black’s laboratory.

### Cell lines and transfection

HEK-293T cells expressing ACE-2 (obtained from BEI resources), A549(ATCC) lung epithelial cells and Vero E6 cells were grown in DMEM media (Gibco) supplemented with 10% FBS (Gibco). For luciferase assays in A549 cells, day before transfection 1.3 × 10^5^ cells were distributed per well of 24 well tissue culture plate. Before transfection DMEM was replaced with Opti-MEM I reduced serum media (Gibco). Lipofectamine 2000 (Invitrogen) was used for all transfection experiment as per manufacturer’s protocol. To check the effect of HuR overexpression, we co-transfected 250ng of pGL3-SARS-CoV-2 5’UTR with either 250ng of pcDNA3.1 or pCDNA3.1-HuR. pGL3 control vector was used for control set. 2ng pRLTK was also co-transfected in each well to normalize the firefly luciferase activity. For HuR knockdown experiment we transfected 150nM of either DsiHuR: 5’AUUUCUGAAUCUGUGACGCAAGAAT 3’(IDT) or Nonspecific (Eurogentec) siRNA 24 hrs prior to luciferase construct transfection.

### Virus stock preparation and infection

Virus stocks were procured from BEI resources, NIAID, NIH and maintained by Viral BSL3 repository at IISc. WT virus: Isolate Hong Kong/VM20001061/2020, NR-52282; Alpha variant (Lineage B.1.1.7): Isolate hCoV19/England/204820464/2020, NR-54000; Beta variant (Lineage B.1.351): Isolate hCoV-19/USA/MD-HP01542/2021,NR-55282; Gamma variant (Lineage P.1): Isolate hCoV-19/Japan/TY7-503/2021, NR-54982; Delta variant (Lineage B.1.617.2): Isolate hCoV-19/USA/PHC658/2021, NR-55611. Vera E6 cells were used to propagate and titrate all the viruses as described before[36]. For virus experiments, HEK-293T-ACE-2 cells were seeded in 24-well plates. The cells were transfected with 150 nM of Nsp si or si HuR using Lipofectamine 2000 as per the manufacturer’s protocol. 16h after transfection, cells were infected with virus stock at an MOI of 0.1 or 1 as described.

### Dual luciferase assay

Luciferase assay was done by dual luciferase assay kit (Promega) according to manufacturer’s protocol. For HuR overexpression and siHuR experiment, assays were done 12 and 8 hrs post-transfection of luciferase constructs, respectively. Luciferase reading was taken with Glomax Luminometer (Promega).

### Protein purification

*E. coli BL21* cells were transformed with pET28a containing His tagged HuR. Recombinant protein was purified with Ni-NTA agarose. The detail of purification was described earlier. Briefly, the expression of HuR was induced at an optical density of 0.6 at 600 nm with 0.5 mM isopropyl-1-thio-β-D-galactopyranoside (IPTG) for 4h. The cells were pelleted and resuspended in lysis buffer (50 mM Tris, pH 7.5, 300 mM NaCl, 0.1 mM phenylmethylsulfonyl fluoride [PMSF]) containing 1X Bacterial protease Inhibitor (Sigma) and lysed by sonication on ice. The supernatant obtained thereafter was incubated with Ni-NTA slurry for 3 h and bound protein was eluted with 500 mM Imidazole after washing the beads with lysis buffer containing 40 mM imidazole. The eluted protein was dialysed in 10 times volume dialysis buffer (25 mM Tris, pH 7.4, 100 mM KCl, 7 mM β-mercaptoethanol [β-ME], 10% glycerol), aliquoted and stored in −70°C.

### *In vitro* transcription

The pcDNA3.1-SARS-CoV-2 5’UTR was linearized with HindIII digestion and purified with phenol-chloroform method. The linearized DNA was used as a template for synthesis of RNA using T7 RNA Polymerase (ThermoScientific) and ^32^P uridine triphosphate. The transcription reactions were carried out under standard conditions at 37 °C for 1.5 h. After alcohol precipitation, RNA was resuspended in nuclease free water. For radiolabelled RNA, 1 μl the prepared RNA was spotted onto DE81 filter paper, washed with phosphate buffer and dried and incorporated radioactivity was measured using scintillation counter.

### UV induced crosslink of RNA and protein

UV-induced cross-linking was carried out as described previously. Briefly, - ^32^P-labeled RNA probes were allowed to form complexes with recombinant HuR in RNA binding buffer (5 mM HEPES, pH 7.6, 25 mM KCl, 2 mM MgCl2, 3.8% glycerol, 2 mM dithiothreitol [DTT], and 0.1 mM EDTA) and then UV irradiated at 254nm for 20 min. The mixture was treated with 50 µg of RNase A (Sigma), separated on an SDS–10% polyacrylamide gel (SDS-PAGE), and band analyzed by phosphorimaging. The 1-300 nts of SARS-CoV-2 genome were used as SARS-CoV-2 5’UTR to protect the secondary structures in the UTR. The multiple cloning site of pGEX vector was transcribed and used as non-specific RNA.

### Surface Plasmon Resonance

Surface plasmon resonance spectroscopy was performed using a BIAcore3000 optical biosensor (GE Healthcare Lifescience) to study the binding kinetics of HuR with SARS CoV2 5’UTR RNA. Biotin-labeled SARS CoV2 5’UTR RNA was immobilized on streptavidin-coated sensor chips (GE Healthcare Lifescience) to a final concentration of 300 Resonance Units (RU)/flow cell. RNA–protein interactions were carried out in a continuous flow of Tris buffer (25 mM Tris (pH 7.5), 100 mM KCl, 7 mM β-mercaptoethanol, and 10% glycerol) at 25 °C at a flow rate of 10 µl/min. Increasing concentrations of HuR protein loaded on the biosensor chip for 100 s (characterized as the association phase), followed by a dissociation phase of 300 s with buffer alone. For normalising background non-specific interaction, a blank surface without any RNA was used for simultaneous injections of the sample during the experiment. BIAevaluation software (version 3.0) was used to determine the on rate, *k*on (M^-1^ s^-1^), and off rate, *k*off (s^-1^), using a 1:1 Langmuir binding model. The binding affinity, *Kd* was determined using the following equation: *Kd* =*k*off/*k*on.

### Plaque forming assay

Plaque assay was performed in Vero E6 cells in 12-well plates. 0.5*10^6^ cells were seeded to give 90% confluency. After 12-16 h, cells were incubated with 100 µl of increasing dilution of supernatant obtained from Nsp si or si HuR treated ACE2-HEK-293T cell infected with SARS-CoV2 or supernatant from mock infected cells. After 1 h of incubation with the cells at 37°C, viral inoculum was removed and cells overlayed with DMEM containing 0.8% Agarose and kept at 37°C. After 48 h of incubation, cells were fixed with 4% paraformaldehyde for 1 h and stained with Crystal violet for 20 mins. Overlay was removed and the number of plaques counted. Following calculation was used to determine PFU/ml: PFU/ml= No. of plaques --; (Dilution factor x Vol of virus inoculum in ml)

### Immunofluorescence

For immunofluorescence staining, HEK-293T-ACE-2 cells were seeded in a 24-well plate on coverslips coated with poly-L-lysine for 14-16 h followed by infection in SARS-CoV2 virus at an MOI of 0.1. After desired time of infection, cells were washed twice with 1X PBS and fixed using 4 % formaldehyde at room temperature for 20 min. After permeabilization by 0.1 % Triton X-100 for 5 min at room temperature, cells were incubated with 3 % BSA at 37 °C for 1 h followed by incubation with the indicated antibody for 2 h at 4 °C and then detected by Alexa-633-conjugated anti-mouse or Alexa-488 conjugated anti-rabbit secondary antibody for 30 min (Invitrogen). Images were taken using Zeiss microscope and image analysis was done using the Zeiss LSM or ZEN software tools.

### HuR knockout cell line preparation

The guide RNA targeting HuR sequence were designed using Zhang lab software. These gRNAs were cloned in the pSpCas9(BB)-2A-GFP (PX458) vector as per the described protocol. The clones were transfected in HEK-293T-ACE-2 cells and 48h post transfection, the cells expressing GFP were sorted as one cell per well in a 96-well plate. These cells were grown and analysed for HuR knockout using western blotting and confirmed though sequencing. For control, untransfected cells were also sorted as one cell per well in parallel and one of the clones, expressing ACE-2 receptor was used.

gRNA1: ACCACATGGCCGAAGACTGC

gRNA2: CTTATTCGGGATAAAGTAGC

### MTT assay

Proliferation was measured using 3-(4,5-dimethylthiazol-2-yl)-2,5-diphenyltetrazolium bromide (MTT). MTT was added to the cells at a final concentration of 0.5mg/ml at the desired time points. After 3-4 h, media was removed, cells treated with 100µl DMSO and the absorbance measured at 560 nm.

### IP-RT

HEK-293T-ACE-2 cells were infected with the SARS-CoV-2 virus at an MOI of 0.1 and after completion of the experiment, 1% formaldehyde was added to the media and incubated at 4 °C for 10 mins. Thereafter, the reaction was quenched with 0.1 M Glycine incubated at 4 °C for 10 mins. The cells were then washed with PBS and lysed in polysome lysis buffer (100 mM KCl, 10 mM HEPES pH 7.0, 5 mM MgCl_2_, 0.5 % NP-40, 1 mM DTT, 100 U/ml RNasin). The supernatant obtained was precleared with Protein G Sepharose beads. In parallel, the Protein G Sepharose beads were incubated with 1 ug of HuR/ PTB antibody overnight at 4 °C and added to the precleared lysates followed by overnight incubation with continuous mixing on a rotator device at 4°C. The beads were then washed three times with polysome lysis buffer. SDS sample buffer was added to the 20% beads and boiled to release the immunoprecipitated protein, and the supernatant was electrophoresed on SDS-12 % PAGE. The RNA was isolated from remaining 80% beads. The beads were incubated with 0.1% SDS and 30 µg proteinase K at 50 °C for 30 mins and RNA isolated after addition of 3 times volume of Tri reagent.

### RNA isolation followed by real time PCR

The total RNA from ACE-2-HEK-293T cells was isolated using TRI Reagent (Sigma) as per manufacturer’s protocol. cDNA was synthesized using following primers: CoV2 Fwd: 5’ TGTCGTTGACAGGACACGAG 3’, CoV2 Rev: 5’ TTACCTTTCGGTCACACCCG 3’, GAPDH Fwd: 5’ CAGCCTCAAGATCATCAGCAAT 3’, GAPDH Rev: 5’ GGTCATGAGTCCTTCCACGA 3’. For quantification of viral positive strand, CoV2 Rev and GAPDH Rev primers were used for cDNA synthesis. For quantification of viral negative strand, CoV-2 Fwd and GAPDH Rev primers were used for cDNA synthesis. Moloney murine leukemia virus(M-MLV) reverse transcriptase was used for cDNA synthesis at 42°C for 1 h using 600 ng of total RNA. Quantitative RT-PCR (qRT-PCR) was done using DyNAmo HS SYBR green qPCR kit (ThermoScientific). qRT-PCR was done using 2 µl of cDNA in a 10-µl reaction mixture according to the manufacturer’s instructions for 40 cycles. The comparative threshold cycle (CT) method was used to calculate fold change in SARS CoV2 RNA levels (2^ΔΔCT) and normalized to GAPDH.

### Western blotting

Protein concentrations of the extracts were assayed by Bradford reagent (Bio-Rad) and equal amounts of cell extracts were separated by SDS-10 % PAGE and transferred onto a nitrocellulose membrane (Sigma). Samples were then analyzed by western blot using the desired antibodies, anti-SARS-CoV-2 N protein (40143-MM05, Sino Biological), anti-HuR antibody (3A2, Santa Cruz), anti-Myc tag antibody (ab9106) or anti-ACE-2 antibody (ab15348) followed by the respective secondary antibodies (horseradish peroxidase-conjugated anti-mouse or anti-rabbit IgG; Sigma). Mouse-monoclonal anti-β-actin-peroxidase antibody (A3854, Sigma) was used as a control for equal loading of total cell extracts. Antibody complexes were detected using the ImmobilonTM Western systems (Millipore).

### Sucrose gradient polysome fractionation

After completion of the experiment, cells were treated with 100 µg/ml cycloheximide for 10min at 37°C. Thereafter, media was removed and cells washed first with ice cold PBS containing 100µg/ml cycloheximide and then with 1X hypotonic buffer (5 mM TRIS-HCl pH-7.5, 5 mM MgCl_2_ and 1.5 mM KCl) containing 100 µg/ml cycloheximide. Cells were scraped in 350µL ice cold lysis buffer (5 mM TRIS-HCl pH-7.5, 5 mM MgCl_2_, 1.5 mM KCl, 100 µg/ml cycloheximide, 1mM DTT, 200 U/ml RNase in from Promega, 0.5% Sodiumdeoxycholate, 0.5% Triton X −100, 200µg t-RNA and 1X protease inhibitor cocktail) and incubated for 15 min on ice. The KCl concentration in the lysate was adjusted to 150 mM. The lysate was spun for 8min at 3000 g at 4°C and supernatant collected. 500 µg of the lysate was loaded on the top of 15–50% sucrose gradient containing 100µg/ml cycloheximide and the gradients were centrifuged at 36,000 rpm for 2 h at 4°C in SW41 rotor (Beckman). Density Gradient Fractionation System (ISCO) was used to fractionate the gradients at a flow rate of 0.3 mm/sec with the UV detector sensitivity set at 1.0. Monosome and Polysomes fractions were pooled to isolate RNA using Trizol (Sigma). SARS-CoV-2 RNA level was quantified using real time PCR.

### Primers for PCR

Genomic 5’UTR Forward: TGTCGTTGACAGGACACGAG

Genomic 5’UTR Rev: TTACCTTTCGGTCACACCCG

N sgmRNA LS Forward: CGATCTCTTGTAGATCTGTTC

N sgmRNA Rev: AGCGGTGAACCAAGACGCA

S sgmRNA LS Forward: CCAACTTTCGATCTCTTGTAG

S sgmRNA Rev: AGAACAAGTCCTGAGTTGAATG

IL-6 Fwd: GGTACATCCTCGACGGCATCT IL-6

Rev: GTGCCTCTTTGCTGCTTTCAC

### Primers for HuR gene PCR

Forward primer: TACTTGCTCTTTTTCTCTTGGC

Reverse primer: CACAGCCATCGTTTCAAGGC

### Anti-sense oligos

Ctrl ASO: CGTTAGATTACCGCG

ASO5: CAACACGGACGAAACC

#### Softwares

All emerging lineage of SARS-CoV-2 were downloaded from Global Initiative on Sharing All Influenza Data (GISAID, https://www.gisaid.org/). For Multiple sequence alignment Clastal Omega online tool (https://www.ebi.ac.uk/Tools/msa/clustalo/) was used. To predict the secondary structure RNAstructure online tool was used (https://rna.urmc. rochester.edu/RNAstructureWeb/Servers/). RNA binding protein was predicted with RNA Binding Protein Database (RBPDB, http://rbpdb.ccbr.utoronto.ca/). Binding region on the RNA sequence was predicted with Cat-Rapid fragment (http://service.tartaglialab.com/page/catrapid_group).

Data was plotted using GraphPad Prism.

## Acknowledgements

SD acknowledges the J.C. Bose Fellowship from SERB, Department of Science and Technology (DST), India, for research support. This study was also supported by Department of Biotechnology (DBT), Government of India, Indo-Swiss project, DBT-IISc partnership program, DST Fund for Improvement of Science and Technology Infrastructure (DST-FIST) level II infrastructure, and the University Grants Commission Centre of Advanced Studies. ST acknowledges financial support from DBT-BIRAC(BT/CS0007/CS/02/20) and Crypto Relief Funds (SP/CRRE-21-0004.05). HR is supported by the research fellowship from the Council of Scientific and Industrial Research (CSIR-SPM). We recognize the SPR facility at the Biological Science division at IISc. We acknowledge BEI resources, NIAID, NIH for providing the SARS-CoV-2 virus strains. We acknowledge Prof. Adolfo Garcia-Sastre, Icahn School of Medicine, for his insightful inputs. We acknowledge viral BSL3 facility at CIDR, IISc supported by DBT-BIRAC and Crypto-Relief foundation for maintaining SARS-CoV-2 virus stocks. We would like to thank Editage (www.editage.com) for English language editing.

---

**Fig S1. Prediction of HuR binding to SARS-CoV-2 5’UTR.** (A) Bioinformatics prediction for binding of RNA binding proteins to SARS-CoV-2 5’UTR using RBPDB. (B) Secondary structure of CoV-2 5’UTR. Red boxes denote the binding sites as found in (A).

**Fig S2. Influence of HuR and cellular and viral functions.** (A) Equal number of WT and HUR KO cells were seeded and their viability of was examined at the described time points using MTT assay (n=5). (B). Effect of HuR overexpression on different SARS-CoV-2 VoCs. HuR KO HEK-293T-ACE2 cells were transfected with either vector control (Control) or HuR overexpression construct (HuR o/e). 16 hrs post transfection cells were infected with different SARS-CoV-2 VoCs at 0.1 MOI. 24 and 48 hrs post infection cells were harvested and viral RNA quantified using specific primers (n=3). Student t-test was used for statistical analysis. *=p<0.05, **=p<0.01, ***=p<0.001.

**Fig S3. Predicted HuR binding sites on SARS-CoV-2 5’UTR.** (A, B) Binding of HuR to SARS-CoV-2 5’UTR using Catrapid software. (C) Sequence alignment of representative 5’UTRs of beta coronaviruses (MERS, SARS-CoV and SARS-CoV-2). The highlighted boxes indicate the shortlisted HuR binding sites.

**Fig S4. The role of HuR in translation.** (A) WT HuR and HuR KO HEK-293T-ACE2 cells were transfected with either pGL3-SV40-Luc (Luc alone) or pGL3-SV40-SARS-CoV-2 5’UTR-Luc (CoV-2 Luc), and RNA levels were quantified 24 hrs post-transfection. (n=3). (B) Area under the curve (AUC) was calculated separately for monosomal peak and the polysomes in uninfected mock profiles of Fig 4D, E from WT and HuR KO cells and the ratio was plotted individually for each set of experiment (n=3). (C) Area under the curve (AUC) was calculated separately for monosomal peak and the polysomes in infected profiles of Fig 4D, E from WT and HuR KO cells and the ratio was plotted individually for each set of experiment (n=3). (D) RNA was isolated from monosome and polysomal fractions (separately) of lysates from SARS-CoV-2 infected WT cells (Fig 4D) and the abundance of Actin and SARS-CoV-2 5’UTR quantified by specific primers in real time PCR. Polysomal loading was calculated as the ratio of RNA in Polysome:Monosomes (n=3). Student t-test was used for statistical analysis. *=p<0.05, **=p<0.01, ***=p<0.001.

**Fig S5. HuR and genomic v/s subgenomic translation. (A)** WT HuR and HuR KO HEK-293T-ACE2 cells were infected with SARS-CoV-2 virus at MOI of 1.0 and 24 h post infection, protein levels of Nsp1, Nsp5 and Nucleocapsid were determined by western blotting using specific antibodies. Ponceau staining of the bot represents equal loading (n=3). **(B)** WT HuR and HuR KO HEK-293T-ACE2 cells were transfected with either *in vitro* transcribed capped Luciferase RNA (Luc alone) or *in vitro* transcribed capped sub-genomic 5’UTR followed by Luciferase RNA (sg 5’UTR luc), and RNA levels were quantified 8h post transfection. (n=3). Student t-test was used for statistical analysis. *=p<0.05, **=p<0.01, ***=p<0.001.

**Fig S6. HuR does not relocalize of cytoplasm upon SARS-CoV-2 infection.** (A)HEK-293T-ACE-2 cells were infected SARS-CoV-2 virus at an MOI of 0.1. Cells were harvested at indicated timepoints and immunofluorescence staining performed using Alexa Fluor conjugated secondary antibodies against HuR (Red) and SARS-CoV-2 Spike protein (Green). The nucleus was counterstained with DAPI. Scale bar represents 10µm. Nuclear-cytoplasmic fractionation was performed at (B) 24 hrs, (C) 48 hrs post infection. Western blotting was performed using anti-HuR antibody to detect the presence of HuR. GAPDH served as cytoplasmic marker and Lamin as nuclear marker.

