## Supplementary figures and images for "HuR enhances SARS-CoV-2 non-structural protein translation through the genomic 5’-UTR, by promoting polypyrimidine tract-binding protein binding"

### Supplementary Fig S1

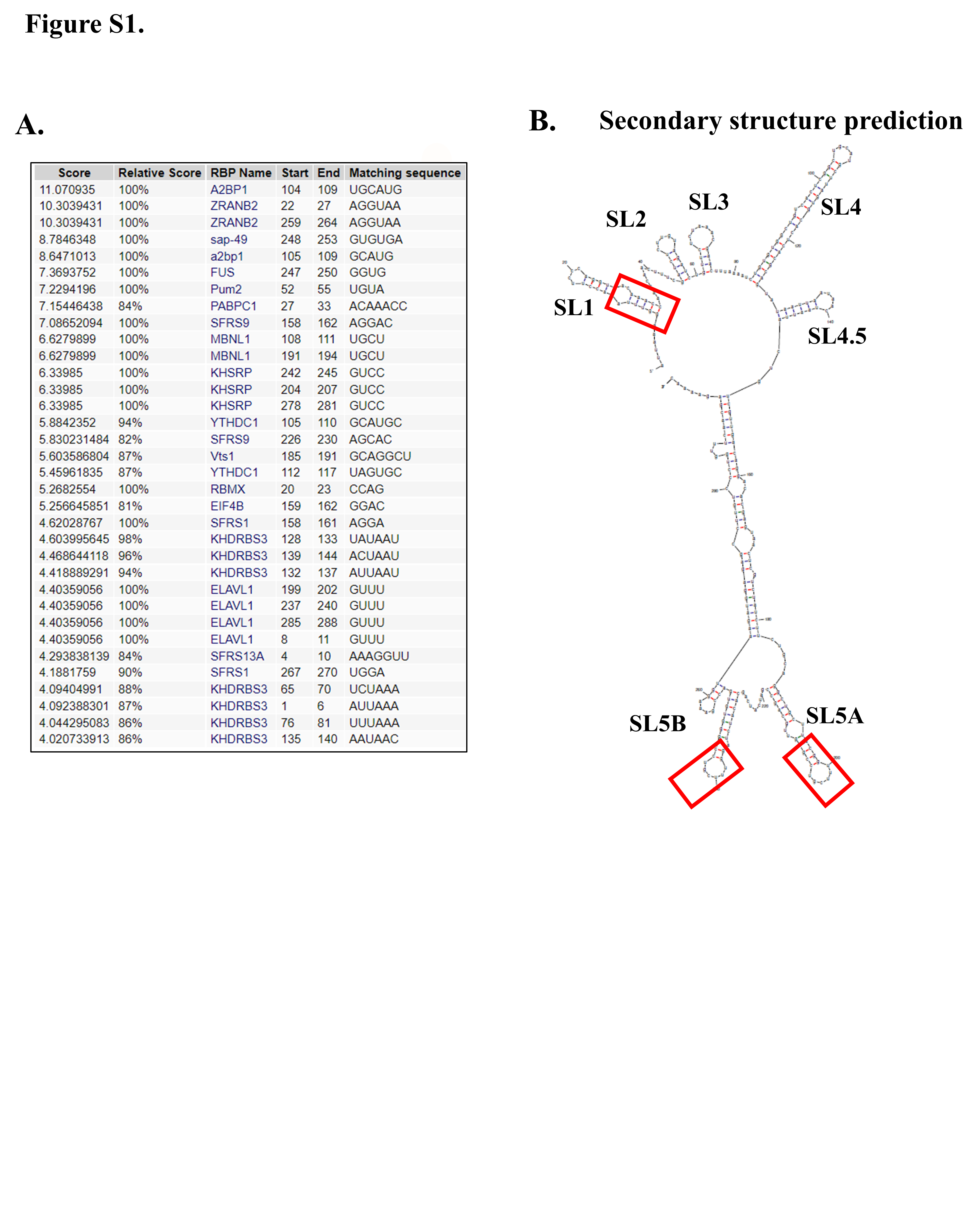

### Supplementary Fig S2

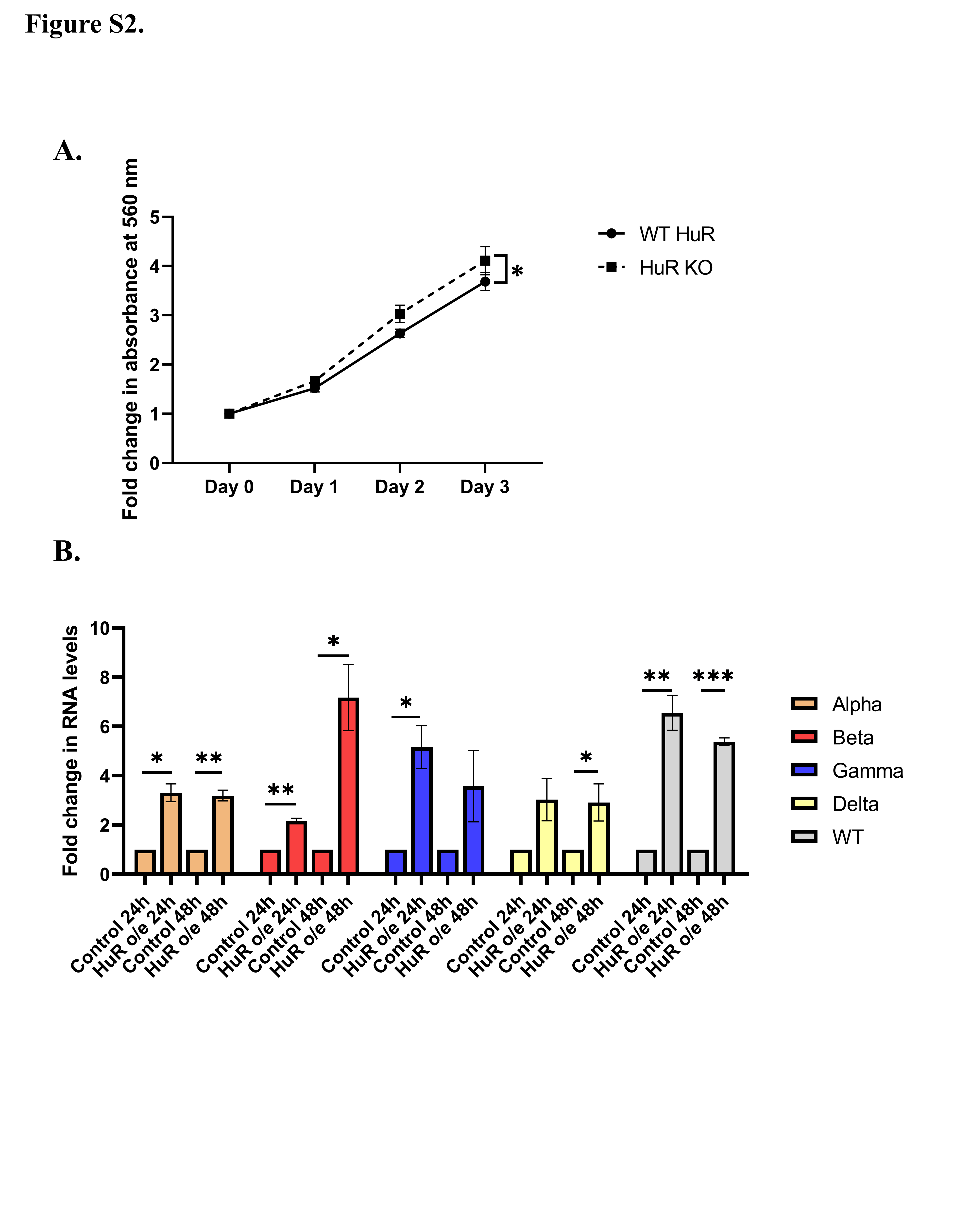

### Supplementary Fig S3

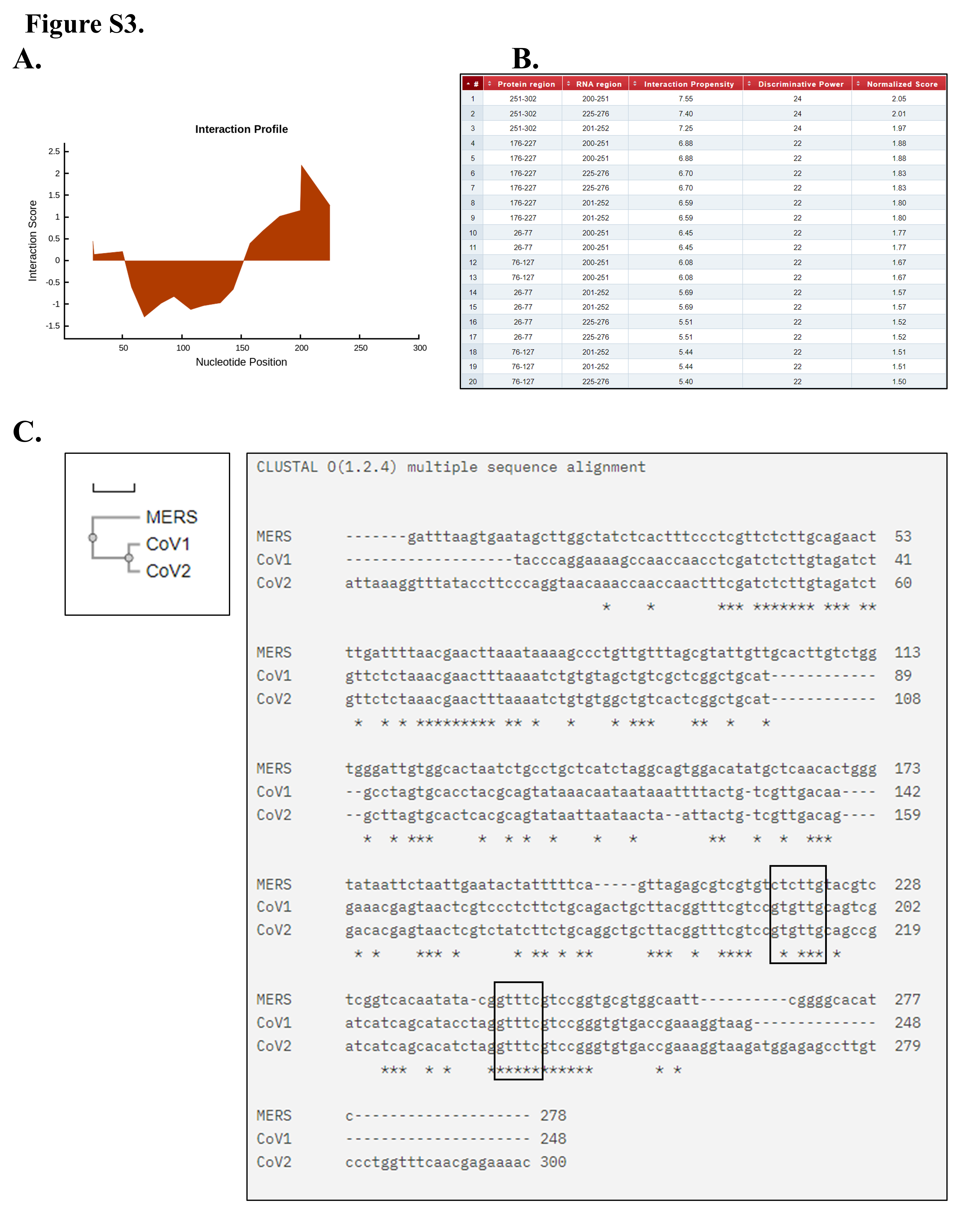

### Supplementary Fig S4

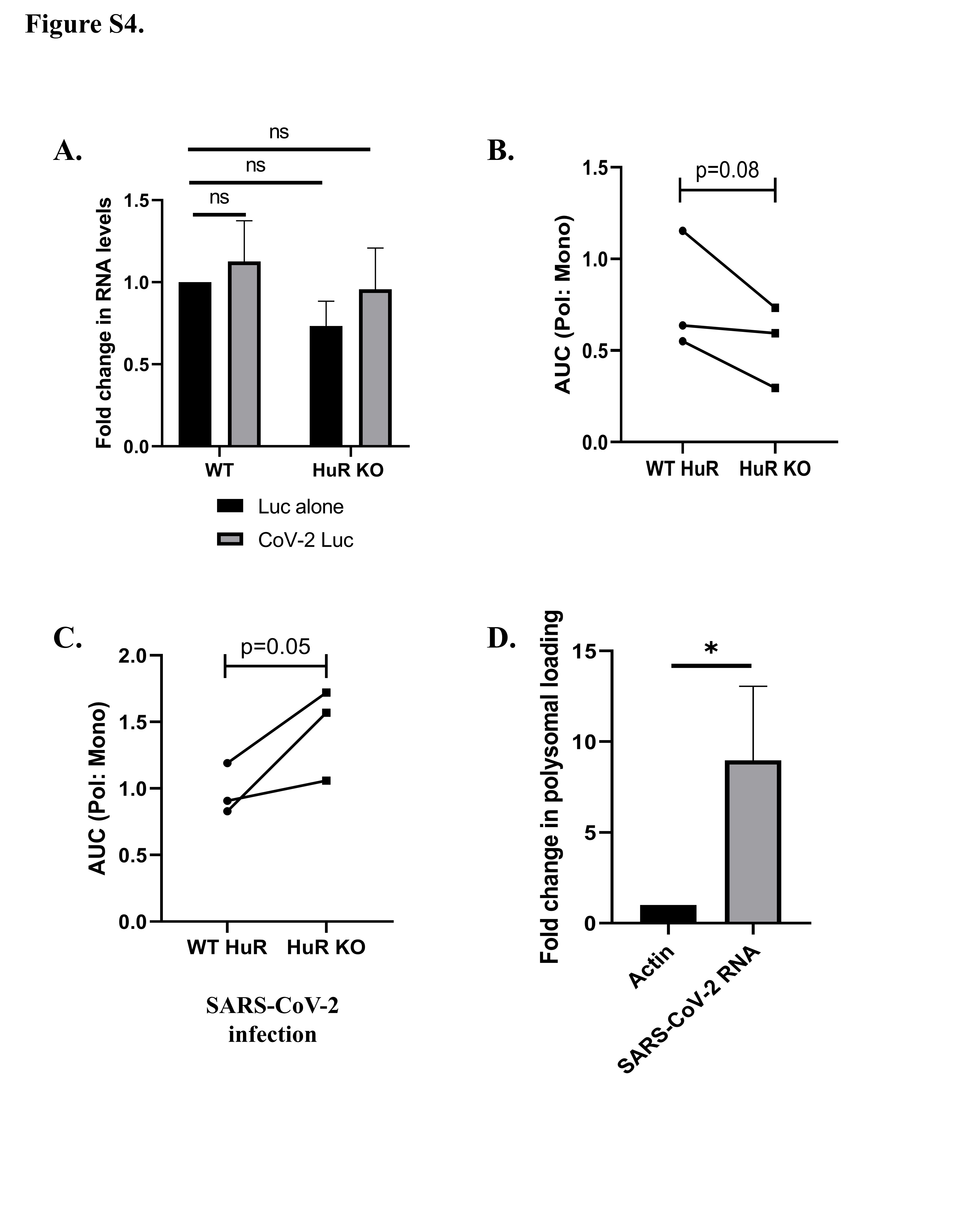

### Supplementary Fig S5

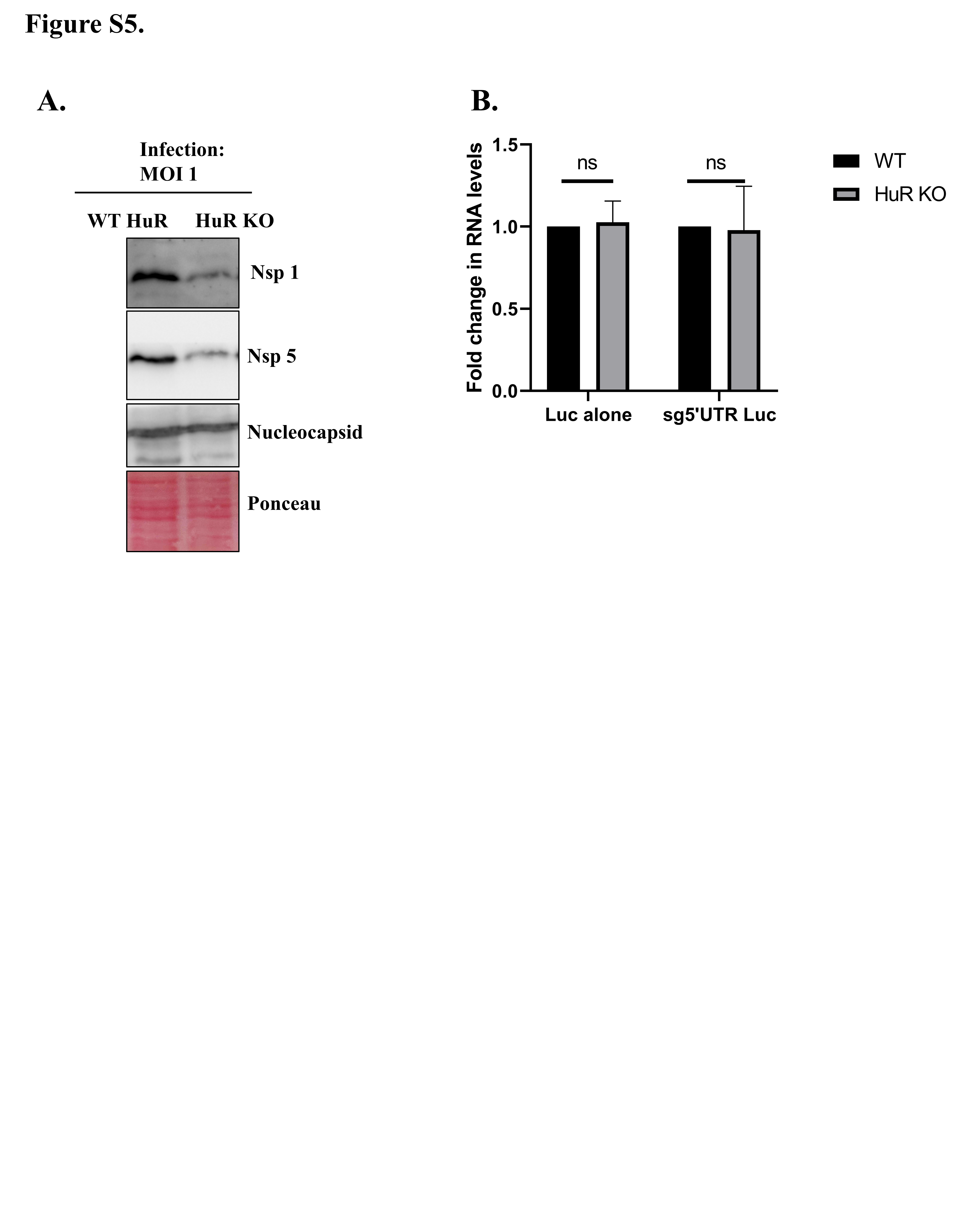

### Supplementary Fig S6

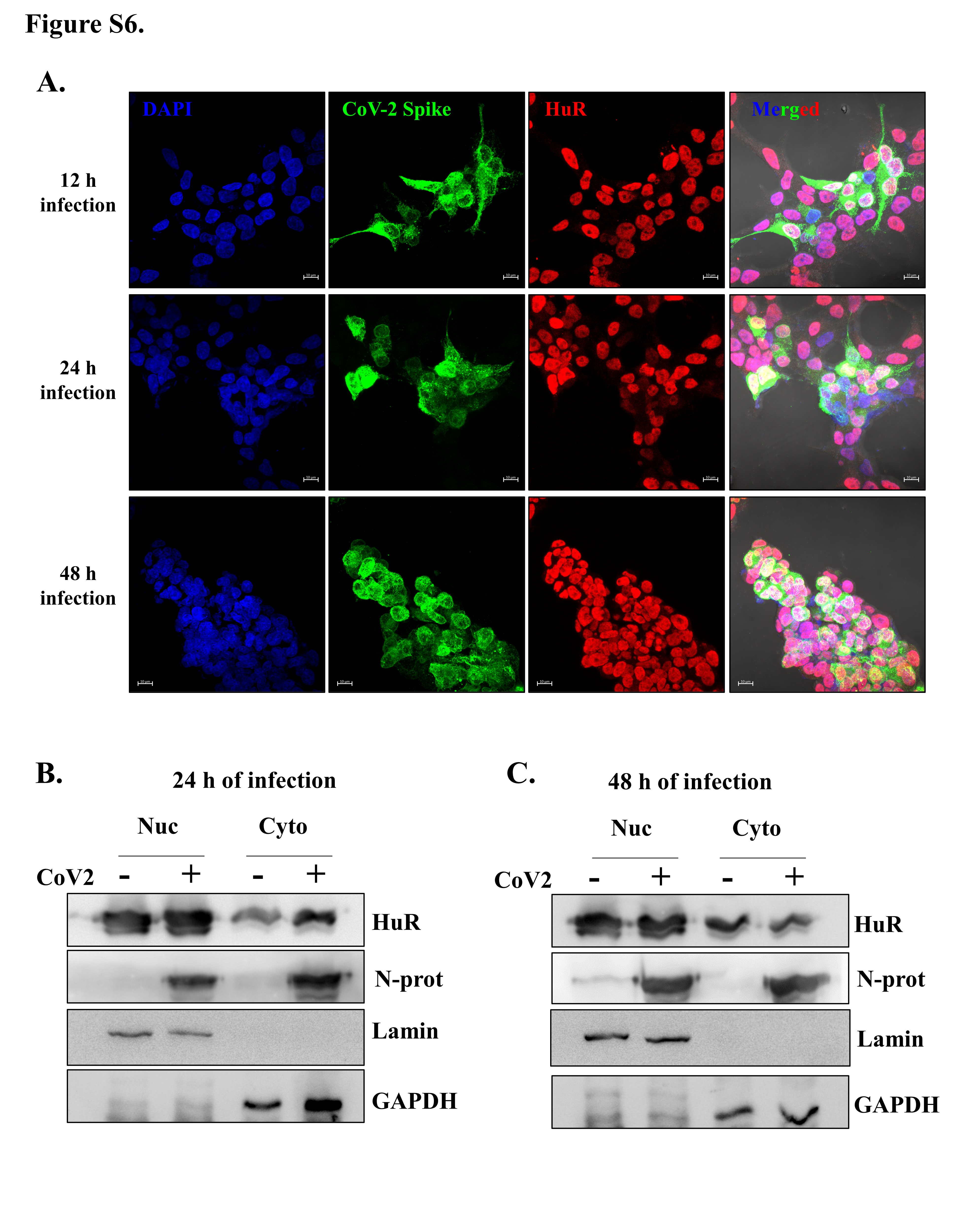
